# A switch in arousal circuit architecture shapes sleep across the lifespan

**DOI:** 10.1101/2025.11.05.686742

**Authors:** Milan Szuperak, Mareike Selcho, Katharina Eichler, Patrick D. McClanahan, Kenta Asahina, Jeffrey B. Rosa, Christopher Fang-Yen, Andreas S. Thum, Matthew S. Kayser

## Abstract

Sleep is a continuous behavior across the lifespan, yet its features and functions evolve markedly with development^1–4^. In *Drosophila melanogaster,* as in mammals, early life sleep differs from mature sleep^5,6^, but it is unknown whether disparate sleep regulatory mechanisms underlie these changes. Here, we identify distinct populations of octopaminergic (OA) neurons that promote arousal in larval and adult flies, thus revealing a developmental switch in sleep-wake circuit architecture. Of eight OA neurons present in the sub-esophageal zone (SEZ) of the nervous system at both life stages, dedicated, non-overlapping subsets drive arousal in larvae versus adults. Morphologic and connectomic analyses show that larval OA arousal neurons project primarily to the ventral nerve cord and lack substantial sensory input, suggesting a circuit logic optimized for internally driven arousal during early development. In contrast, adult OA arousal neurons target higher brain regions involved in cognition and receive rich multimodal sensory input, supporting wakefulness in response to environmental cues. These findings highlight a developmental transition in arousal circuitry that mirrors changing ecological demands, with juvenile systems organized to prioritize growth and feeding, insulated from sensory disturbance, and mature systems supporting sensory-guided behavior. Our results support a model of sleep regulation as a developmentally dynamic process, in which shared neuromodulators like OA operate through distinct cellular substrates tailored to life stage– specific behavioral priorities.

## Introduction

Across the lifespan, numerous sleep properties show dynamic change. In early life, sleep duration and depth are greatest, and as animals age, sleep becomes more fragmented^1,3,4^. Perhaps most notably, REM sleep in mammals is most abundant during infancy before tapering sharply by adulthood^1^. Yet, not all components of REM are present in these earliest stages. In infant rats, for example, limb twitches and muscle atonia are evident in the earliest postnatal period, while cortical EEG signatures of REM do not appear for another 10 days^7^. Similarly, young adult fruit flies sleep more deeply than mature flies, but this juvenile sleep state has behavioral features that are distinct from sleep experienced by mature flies, even following sleep deprivation^8,9^. For example, excess sleep in the juvenile period is driven by reduced probability of transitioning to wake from a sleep state, while a sleep-deprived adult fly exhibits a prominent increase in the probability of falling asleep^8^. Although sleep is thought to be a continuous behavior across the lifespan^10^, it is unknown whether underlying regulatory mechanisms of juvenile and mature sleep are the same.

The ontogenetic hypothesis of sleep postulates that early developmental sleep is important for brain patterning and function^1^. Human and rodent studies have demonstrated that impaired sleep during critical periods of development has severe neurobehavioral consequences^11–16^, and sleep deprivation in young flies impairs adult social behaviors^6,17^. Accumulating evidence suggests that juvenile sleep has distinct genetic underpinnings in comparison to sleep in maturity^18,19^. While numerous genes have been described that control sleep in mature adult flies^20–23^, mutation to most of those same genes has no effect on sleep maturation or sleep during earlier developmental periods^9^. In humans, adult sleep duration-associated loci show little association within childhood/adolescent Genome Wide Associate Studies (GWAS) for sleep duration^18^. While major strides have been made over the past two decades in understanding how neural circuits in the mature brain give rise to sleep^24,25^, little is known regarding sleep regulatory circuits during early phases of life.

As in humans, sleep in fruit flies changes with age: young flies immediately following eclosion exhibit increased sleep amount and depth in comparison to mature adults (∼5-7 days old)^5,6^. Our group also described sleep in larval stages^26^, providing a platform for investigation of sleep during even earlier developmental stages. In larvae, despite the presence of dopaminergic (DA) neurons and their known involvement in memory and other processes at this stage, DA does not appear to be a driver of arousal^26^. Instead, octopamine (OA; insect functional homolog of noradrenaline) is the primary arousal cue, a role it maintains into adulthood^26,27^, by which time DA has “come online” as a wake-promoting cue as well^22,28,29^. These juvenile vs adult distinctions could be consistent with a model in which sleep regulatory mechanisms are elaborated over time, similar to REM sleep, such that core features are present at the earliest stages (OA) while others (DA) are layered on across developmental periods. Alternatively, the regulatory boundaries might be more demarcated, with dissociable mechanisms (e.g., distinct neuronal populations) underlying the seemingly continuous behavioral sleep maturation process.

Here, we use *Drosophila* to investigate the neuronal circuit logic of wakefulness across development. We pinpoint a population of ∼8 OA neurons in the subesophageal zone (SEZ) of the *Drosophila* brain whose activation is sufficient to induce arousal from sleep in larval and adult stages. Surprisingly, intersectional genetic and stochastic labeling approaches demonstrate that the specific OA cells within the SEZ relevant for arousal in larvae compared to adults are completely distinct, even though all 8 OA neurons are present at both life stages. Morphologic and connectomic analyses reveal that the adult OA arousal neurons send ascending projections to innervate brain regions involved in sleep and cognitive processes, while receiving multimodal sensory input. In contrast, larval arousal neurons have descending projections to the ventral nerve cord (VNC) and are nearly sensory devoid. Our findings demonstrate unique neural circuit architectures regulating wakefulness across the lifespan, and suggest a circuit strategy designed to achieve sensory disconnectedness of arousal loci in early life.

## Results

### Identification of larval octopaminergic subpopulations promoting wake

We previously showed that OA neurons play a central role in promoting wake in *Drosophila* larvae^26^. The larval nervous system includes ∼80 OA neurons^30,31^, and we sought to determine if a specific subpopulation of these cells is most relevant for regulating sleep/wake during this developmental period. We performed a thermogenetic neural activation screen in second instar larvae (L2) using a collection of GAL4 drivers that putatively express in OA neurons, based on GAL4 insertion in the proximity of enhancers for either of the OA synthesis enzymes Tyramine β hydroxylase (Tbh) or Tyrosine decarboxylase 2 (Tdc2)^32^. We expressed the heat-sensitive cation channel *UAS-TrpA1*^33^ under control of 14 available GAL4s, and identified 3 that produced a wake-promoting effect when the flies were exposed to elevated temperature. Thermogenetic activation with the drivers R57F09, R76H03 or R76H04 resulted in reduced sleep duration, sleep bout number, and sleep bout length compared to genetic control (Fig. 1a-c). Activation of neurons defined by these GAL4s, especially R57F09, also increased larval activity while awake (Fig. 1d), consistent with what we observe during activation of all OA neurons^26^. To determine which OA neurons are labeled by these drivers, we expressed nuclear GFP under control of each GAL4 and examined both central brains and the ventral nerve cord (VNC). R57F09-GAL4 drove GFP expression in nearly all OA neurons (Fig. 1e), while R76H03 and R76H04 did so in many fewer OA cells, primarily in the subesophageal zone (SEZ), but also in many non-OA neurons throughout the brain (Fig. 1f,g).

**Fig. 1.**
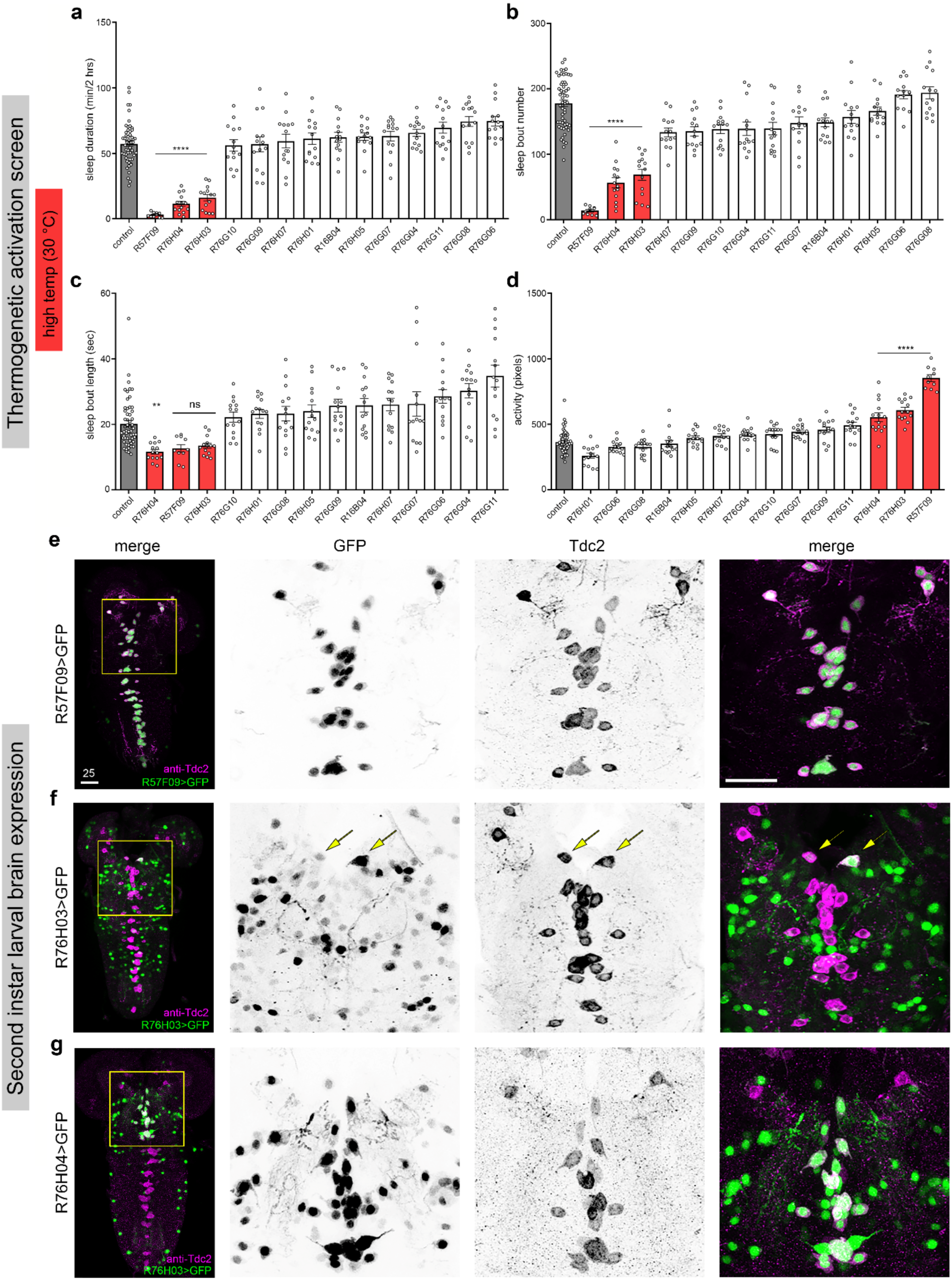
Octopaminergic neurons promote arousal during early *Drosophila* development. **a**, Sleep duration, **b**, sleep bout number, **c**, sleep bout length and **d**, activity values of 2^nd^ instar larvae upon thermogenetic activation of various subsets of OA neurons (n=10-14). All three selected Gal4 drivers are broadly expressed in multiple neurons throughout the 2nd instar larval brain (**e**, **f** and **g**). **e**, *R5709-Gal4* expression is mostly overlapping with all the OA neurons. Panels on the right show zoomed-in-view of the yellow box with individual labeling (GFP for Gal4 expression, Tdc2 for OA neurons) or merge for combined. **f**, *R76H03-Gal4* is expressed only in one pair of OA neurons in the Subesophageal Zone (SEZ) (arrows). **g**, *R76H04-Gal4* expression overlaps with a subset of OA neurons in the SEZ. Images on **e**, **f** and **g** are maximum projections of multiple stacks of the whole brain, the other panels are cropped from the originals. One-way analysis of variance (ANOVA) followed by Tukey’s multiple comparison tests. For this and all other figures unless otherwise specified, data are presented as mean ± SEM; n.s., not significant; *P < 0.05, **P < 0.01, and ***P < 0.001. Scale bars: 25 µm.

We next obtained a split-GAL4^34^ to gain genetic access to larval OA subpopulations of interest in the SEZ. We found that intersection of R76H04-GAL4 and Tdc2-GAL4 (now referred to as OA^SEZ^-split-GAL4) limited expression to a small group of OA SEZ neurons, with the number of labeled cells varying from 4-8 between brains (Fig. 2a-c). Seven of the neurons were already identified^31,35^: six include the symmetrically paired neurons OA-sVPMmx (also known as OAN-g1), OA-sVPMlb, and OA-sVPMmd2, and one was the unpaired neuron OA-sVUMlb3 (Extended Data Fig. 1; Table 1). The final cell is newly identified through single cell labeling approaches in this work and named OA-sVUMlb5. The cell body of sVUMlb5 is in close proximity to sVUMlb3, but the VNC projection patterns of these neurons are distinct, with sVUMlb5 neurites more medial than sVUMlb3 (Extended Data Fig. 1). To directly assess whether OA SEZ neurons promote wakefulness in L2, we performed thermogenetic activation as in the original screen. We observed a reduction in sleep duration and sleep bout length at elevated temperature compared to controls (Fig. 2d-f; Extended Data Fig. 2). We did not detect an increase in larval activity in the experimental group (Fig. 2g), indicating that activation of this OA subpopulation promotes arousal without affecting locomotor behavior while awake. To further assess arousal capabilities of these neurons, we modified existing platforms^36^ to develop a closed-loop optogenetic activation system (Fig. 2h) in which freely-moving larvae in individual arenas were behaviorally-monitored in real time, with a focused optogenetic light stimulus delivered to an animal when quiescence was detected. Using this platform, we observed strong suppression of sleep upon activation of the restricted group of OA neurons compared to controls (Fig. 2i). We noted many instances in the experimental condition where the mean sleep bout length dropped to the minimum possible (6 seconds), meaning activation of OA SEZ neurons immediately elicited arousal (Fig. 2j). The frequent disruption of sleep in the closed loop system was associated with an increase in sleep bout number compared to the controls suggesting greater drive to sleep, but no change in larval activity while awake (Extended Data Fig. 2). Together, these data define a population of 8 OA neurons in the SEZ whose activation is sufficient to induce wakefulness.

**Table 1.**

| larval neuron name | larval neuron driver | adult neuron name | adult neuron driver | additional notes |
| --- | --- | --- | --- | --- |
| sVPMmx/OAN-g1 | SS4268-Gal4 | OA-VPM3 | Tdc2-p65.AD; R76H04-Gal4.DBD<br>MB022B-Gal4 | OA<sup>SEZ</sup>-split-Gal4 (this study) |
| sVPMIb | MB113C-Gal4 | OA-VPM4 | MB021B-Gal4 |  |
| sVPMmd2 | None | OA-VPM2 | Tdc2-p65.AD; R15E12-Gal4.DBD |  |
| sVUMd3 | None | OA-VUMd3 | Tdc2-Gal4;Fru-FLP (observed though not used in this study) | Previous ID VUMd3 #2<sup>42</sup> |
| sVUMd5 | None | OA-VUMd5 | None | Previous ID VUMd3 #1<sup>42</sup> |

**Fig. 2.**
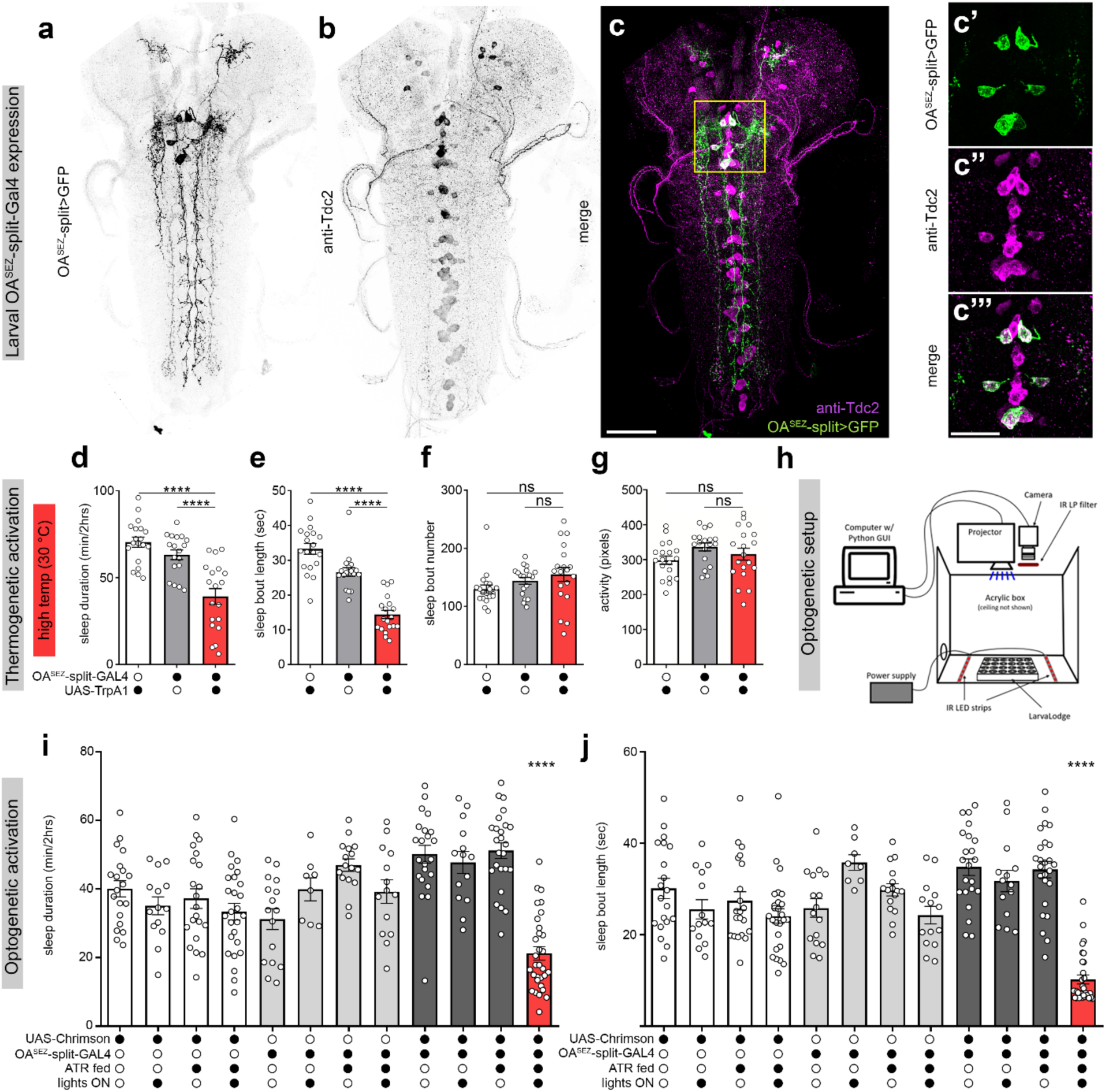
A subpopulation of OANs in the SEZ drive larval wakefulness. **a,** OA^SEZ^-split-GAL4 (intersection of Tdc2-p65.AD and R76H04-Gal4.DBD) labels 4-8 SEZ OANs (GFP) that are Tdc2-positive (**b**). **c,** Merged and higher magnification view of a and b. **d-g,** Thermogenetic activation of OA^SEZ^-split-GAL4 neurons significantly reduced sleep duration and sleep bout without affecting activity of 2nd instar larvae (n=18-19). **h,** Schematic of the closed-loop optogenetic system. **i,j,** Optogenetic activation of OA^SEZ^-split-GAL4 neurons reduced sleep duration and sleep bout length compared to controls (n=8-31). Images on **a**-**c** are maximum projections of multiple stacks of the whole brain. Scale bars: 25 µm.

### Octopaminergic neurons in the SEZ promote wake across the lifespan

OA also promotes arousal in adult *Drosophila*^27,37,38^, but it is unknown if the same circuit logic is shared from larval to adult periods. To address whether OA SEZ neurons maintain a similar function in adulthood, we first examined the adult expression pattern of OA^SEZ^-split-GAL4. As in larvae, GAL4 expression was somewhat stochastic but we typically detected 1-3 distinct OA populations in the adult nervous system. One paired group of OA neurons (OA-VPM3) was always present (Fig. 3a-c; Extended Data Fig. 3). OA-VPM3 are known to survive metamorphosis and correspond to larval OA-sVPMmx^39^, which we characterized as among the larval OA neurons of interest. OA-VUMa2 was also labeled in most adult brains, and we sometimes observed OA-VPM4 (Extended Data Fig. 3). The larval neuron corresponding to OA-VUMa2 (sVUM1) was never labeled with OA^SEZ^-split-GAL4, so we did not characterize the adult neuron further. To determine whether the adult OA SEZ neurons with corresponding larval OANs maintain the same wake-promoting role in adulthood, we used a thermogenetic approach with assessment of sleep in high-resolution multibeam *Drosophila* activity monitors^40^. Compared to baseline sleep measures, activation of these OA SEZ cells in the adult resulted in a dramatic reduction in sleep duration during the night (Fig. 3d-f; Extended Data Fig. 4). These findings suggest that the OA neurons we identified maintain their arousal-promoting role from larval life to adulthood.

**Fig. 3.**
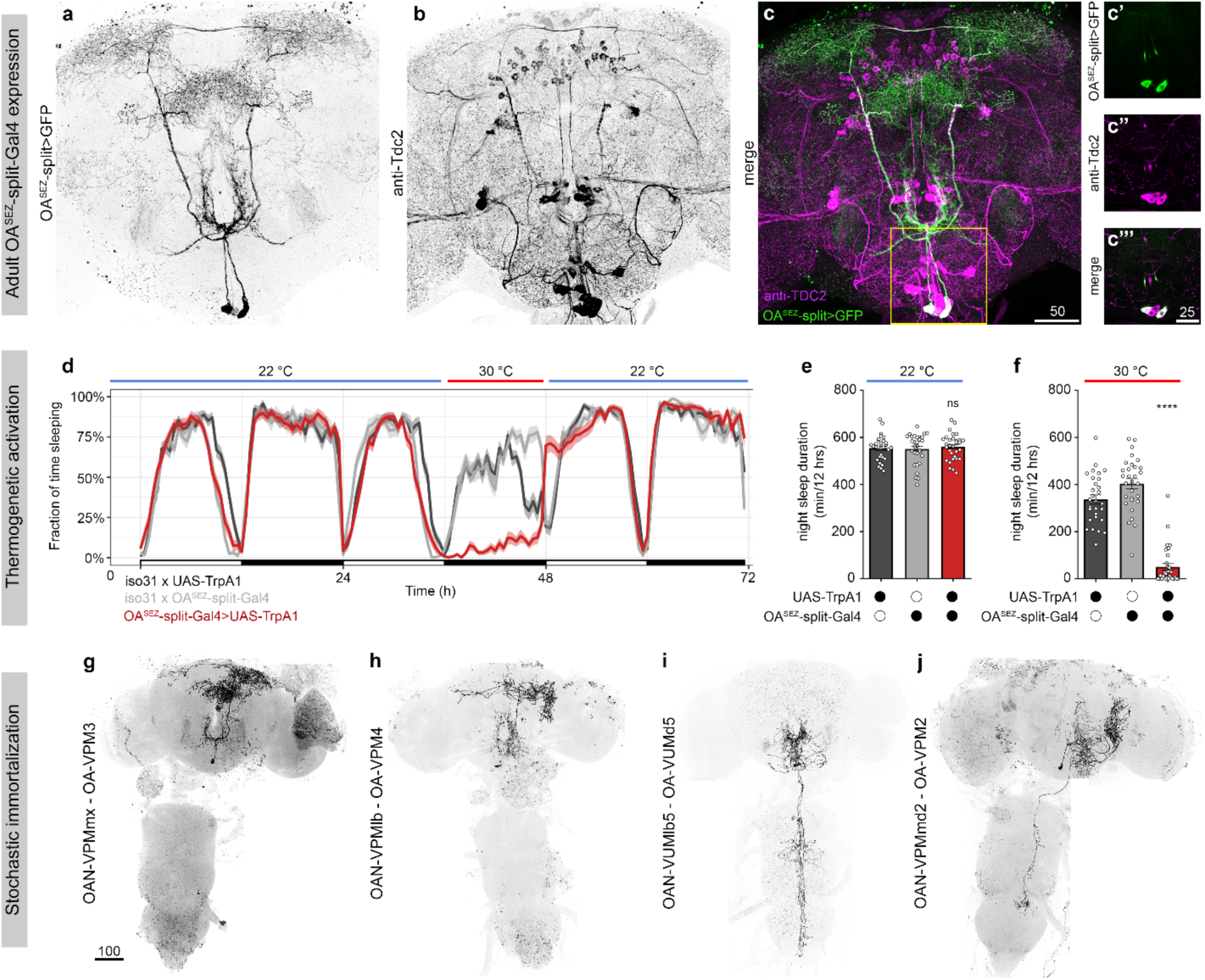
Neurons defined by OA^SEZ^-split-GAL4 survive metamorphosis and promote adult arousal. **a,b,** OA^SEZ^-split-GAL4 (GFP) is expressed in ascending OANs (anti-Tdc2) in the adult brain. Merged and higher magnification in **c**. **d,** Sleep traces over 3 consecutive days demonstrating thermogenetic activation of OA^SEZ^-split-GAL4 neurons reduced nighttime sleep of adult flies. **e,f,** Quantification of night sleep duration at restrictive (22 °C) or permissive (30°C) temperature compared to controls (n=30-32). **g-j,** Immortalized fluorescence in larval OA^SEZ^-split-GAL4 neurons reveals survival of each cell to adulthood. Scale bars: **a**-**c** = 50 µm, insets = 25 µm, and **g**-**j** = 100 µm.

To determine whether these and other of the OA SEZ neurons of interest survive metamorphosis and persist into adulthood, we used a fluorescence immortalization strategy^41^. A hormone-inducible FLP recombinase was expressed using OA^SEZ^-split-GAL4 with FLP activity induced during the second instar larval stage, facilitating permanent expression of *LexAop-Chrimson:Venus* for visualization in adult brains. The labeling of OA SEZ neurons was stochastic, allowing us to determine the origin of individual neurons by comparing larval and adult projection patterns. With this approach, we verified that adult OA-VPM3 neurons are derived from the larval OA-sVPMmx neurons (Fig. 3g). Using a larval OA-sVPMlb-specific driver (MB113C-GAL4), we also demonstrated that adult OA-VPM4 neurons correspond to the larval OA-sVPMlb neurons (Fig. 3h; Extended Data Fig. 5). Surprisingly, although the adult OA^SEZ^-split-GAL4 expression pattern only includes OA-VPM3/4 and VUMa2, we found that additional larval OA^SEZ^-split-GAL4 neurons survive metamorphosis. GAL4 expression patterns are appreciated to change across developmental stages^42^, and tracking the OA^SEZ^-split-GAL4 expression pattern across development indeed revealed disappearance of GFP signal in some subpopulations during pupal stages (Extended Data Fig. 6). In contrast, immortalization strategies demonstrated that the newly-identified larval OA-sVUMlb5 neuron becomes an unpaired adult OAN, here named OA-VUMd5 (Fig. 3i). While OA-VUMd5 exhibits similar brain morphology to a previously characterized OA-VUMd3^43^, innervation patterns within the SEZ and VNC are distinct, as in larvae (Extended Data Fig. 7). We also confirmed that larval OA-sVPMmd2 neurons develop into adult OA-VPM2s (Fig. 3j)^31,44^. Thus, all of the larval OA SEZ neurons of interest are maintained in the adult nervous system (Table 1).

### Distinct individual octopaminergic neurons regulate sleep/wake in larvae vs adults

Given that OA^SEZ^-split-GAL4 labels distinct neurons/pairs, we next investigated whether any of these subpopulations could be pinpointed as sufficient for promoting arousal in larvae or adults. We obtained a split GAL4 driver line, SS4268-GAL4, that specifically labels larval sVPMmx neurons (Fig. 4a), which correspond to adult OA-VPM3. Optogenetic activation of larval sVPMmx or sVPMlb neurons (using MB113C-GAL4) had no effect on sleep duration or sleep bout length (Fig. 4b-f; Extended Data Fig. 8), the key sleep metrics altered with activation of all neurons defined by OA^SEZ^-split-GAL4. Thus, despite a strong arousal-promoting effect of VPM3/4 in adulthood (see Fig. 3), our results demonstrate the corresponding larval neurons are not relevant for sleep/wake state changes. Critically, sVPMmx neurons have a well-known role in larval memory processing^31,45^, arguing against general non-functionality of these cells during this period.

**Fig. 4.**
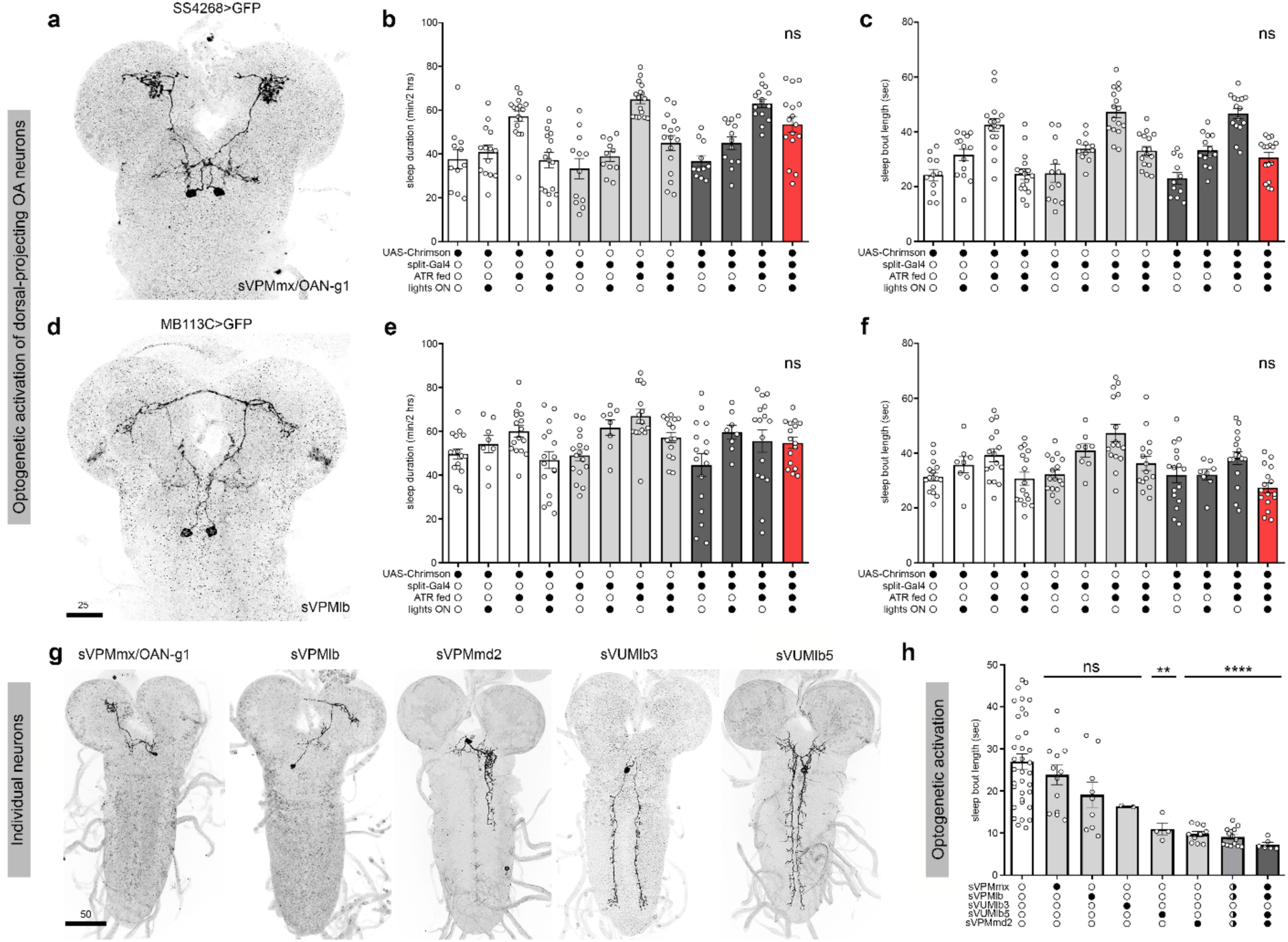
A subpopulation of descending OANs are responsible for larval arousal. **a,** SS4268-split-GAL4 expresses in the larval OA-sVPMmx neurons. **b,c,** Optogenetic activation of OA-sVPMmx does not affect sleep duration or sleep bout length (n=11-16). **d,** MB113C-split-GAL4 expresses in larval OA-sVPMlb neurons. **e,f,** Optogenetic activation of OA-sVPMlb does not affect sleep duration or sleep bout length. (n=8-16). **g,** Visualization of specific larval OAN cell types within the population defined by OA^SEZ^-split-GAL4 when using stochastic labeling approaches. **h,** Sleep bout length with optogenetic activation of individual or various combinations of stochastically labeled OANs. Partially filled circles represent brains where at least one ascending- and one descending-projecting neuron was present simultaneously. Full circles represent animals/brains where at least one of each neuron of interest was included. Each data point represents 1 larva. Scale bars: **a** and **d** = 25 µm, **g** = 50 µm.

Are the remaining untested neurons defined by OA^SEZ^-split-GAL4 sufficient to promote arousal in 2^nd^ instar larvae? Since we were unable to identify specific drivers to gain stable access to these cells, we instead took advantage of stochastic labeling inherent to immortalization strategies. Specifically, immortalized/labeled neurons express *LexAop-Chrimson:Venus*, allowing us to optogenetically activate only those cells in the closed-loop system, assess the wake-promoting effect on a per animal basis, and subsequently dissect brains from individual larvae with visualization of the Venus tag to determine which neurons were activated (see individually immortalized neurons in Fig. 4g). Consistent with results from our stable subpopulation driver lines (Fig. 4a-f), animals in which only sVPMmx and/or sVPMlb neurons were activated had no effect on sleep (Fig. 4h; Extended Data Fig. 9). We only found two cases where the sVUMlb3 neuron was individually immortalized and activated, and we observed that the sleep bouts were not significantly reduced. In contrast, activation of one or both sVPMmd2s, or the unpaired sVUMlb5 neuron, had a strong effect on sleep, arousing the larvae almost immediately (represented by near minimum sleep bout length of 6 seconds) (Fig. 4h; Extended Data Fig. 9). This wake-promoting effect was not strengthened by combinations of these cells or additional stochastic labeling of sVPMmx or sVPMlb, though further reduction of sleep might not be possible due to a floor effect. Notably, sVPMmd2 and sVUMlb5 are descending cells innervating the VNC while the ascending sVPMmx and sVPMlb neurons arborize within the central brain (Fig. 4g), suggesting descending OA neurons have a privileged role in arousal during early life.

We next examined whether specific OA subpopulations are crucial for arousal in adulthood. OA^SEZ^-split-GAL4 labels OA-VPM3, OA-VPM4, and OA-VUMa2, and activation of these neurons indeed promoted wake (Fig. 5a,f-g). Another driver, MB022B, likewise expresses in OA-VPM3 and OA-VPM4 (Fig. 5b) and activation of the cells suppressed night sleep to a similar degree as OA^SEZ^-split-GAL4 (Fig. 5h). While we were not able to identify an OA-VPM3-specific GAL4, we next tested activation of OA-VPM4 (MB021B-GAL4) and observed a reduction of sleep though to a lesser degree than OA-VPM3 and 4 together (Fig. 5c,i). Moreover, recent work^46^ has shown specific activation of OA-VPM3 is indeed strongly wake-promoting, consistent with our results. Our findings therefore suggest that OA-VPM3 represents a strong wake-promoting locus in the adult brain. Next, we tested the role of adult neurons corresponding to the primary larval arousal cells, sVPMmd2 and sVUMlb5. We used a newly identified split-GAL4 (Tdc2Gal4-AD ∩ R15E12-DBD) that labeled OA-VPM2s, corresponding to larval sVPMmd2s (we also see OA-VUMa2 in ∼50% of brains with this split-GAL4) and found activation of these cells failed to elicit wakefulness in adult flies (Fig. 5d,j; Extended Data Fig. 10). No stable driver line exists to access OA-VUMd5, so we once again used a stochastic immortalization strategy (as in Fig. 4h) with optogenetic activation during sleep assays in adulthood followed by examination of labeled cells in each brain. Animals in which OA-VPM3 expressed *Chrimson:Venus* showed a wake-promoting effect in comparison to flies in which no neurons were labeled (Fig. 5k-n), consistent with results using OA-VPM3 driver lines. In contrast, optogenetic activation of animals with OA-VUMd5 labeled exhibited no arousal phenotype, indicating a loss of the larval arousal role (Fig. 5l,m). These data indicate that OA arousal neurons in the adult brain are dissociable from those in the larval brain, and raise the possibility of an anatomical flip across the lifespan from descending to ascending arousal circuit architectures.

**Fig. 5.**
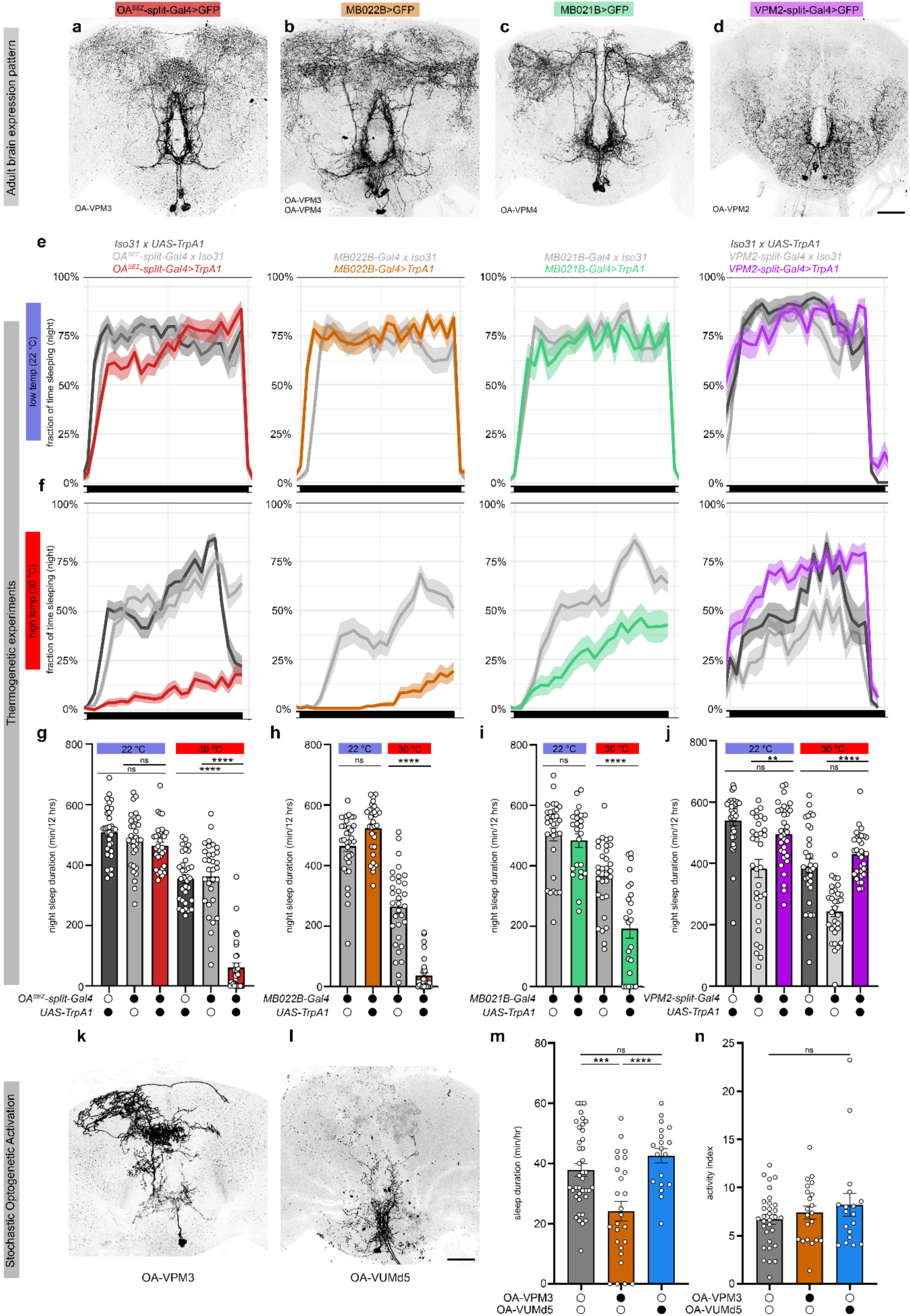
A subpopulation of ascending OANs promotes adult arousal. **a-d,** Expression pattern of split-GAL4 combinations or other intersectional genetic approaches to access SEZ OANs of interest in the adult brain. OA^SEZ^-split-GAL4 labels OA-VPM3, OA-VPM4, and OA-VUMa2 neurons (**a**). MB022B is expressed in OA-VPM3 and OA-VPM4 neurons (**b**). MB021B labels a combination of OA-VPM4 and OA-VUMa2 neurons (**c**). VPM2-split-Gal4 drives expression in OA-VPM2 and OA-VUMa2 neurons (**d**). **e,f,** Nighttime sleep traces at the restrictive temperature (22 °C; **e**) or permissive temperature (30 °C, **f**) during thermogenetic experiments. **g-j,** Sleep duration during the night in each genotype compared to controls. Immortalized fluorescence expression pattern of an OA-VPM3 (**k**) and OA-VUMd5 neuron (**l**). Sleep duration (**m**) and activity (**n**) over 1 hour in control (no neurons labeled; gray), OA-VPM3 (orange), or OA-VUMd5 (blue) labeled animals during optogenetic activation. n=24-32 for **g**-**j**, n=18-31 for **m**-**n**.n=24-32. Scale bar = 50 µm.

### Sensory disconnectedness of arousal circuits in early life

Given the distinct projection patterns of arousal OA neurons in larvae compared to adults, we next investigated potential downstream targets relevant for sleep regulation. While little is known regarding sleep regulation during larval periods, numerous brain regions have been implicated in adult sleep control^25,47^. We first used trans-TANGO^48^ under control of OA^SEZ^-split-GAL4 to visualize postsynaptic targets of OA-VPM3/4 in the adult brain. This approach revealed extensive innervation of the fan-shaped body of the central complex and the mushroom body, two regions known to regulate sleep^49–55^ (Extended Data Fig. 11). Moreover, using the FlyWire adult connectome^56,57^ we found that OA-VPM3 makes monosynaptic connections with sleep-promoting dorsal fan-shaped (dFB) neurons defined by 23E10-GAL4^58^. These OA neurons, therefore, are positioned to promote arousal interaction with known sleep centers. While sleep-promoting neurons have yet to be characterized in larval stages, we note that sVPMmx (no arousal role) directly synapses onto the larval MB acting in a memory capacity^31,45,59^, while sVUMlb5 and sVPMmd2 (arousal-promoting) do not, suggesting that larval arousal OA neurons are unlikely to regulate sleep via MB connectivity.

To further understand arousal circuit organization across developmental periods, we utilized complete larval and adult brain connectomes^57,60–65^, focusing on sensory inputs to OA neurons. In larvae, consistent with OA projection morphologies, we observed that the arousal-irrelevant neurons (sVPMmx or sVPMlb) make nearly all synaptic contacts (inputs and outputs) in the central brain. In contrast, arousal-promoting sVUMlb5 and sVPMmd2 project to the VNC and make most synaptic contacts there. Next, we denoted inputs received by each OA neuron subtype (via an interneuron) from 12 sensory modalities. Strikingly, while sVPMmx and sVPMlb receive input from 7 and 5 sensory modalities, respectively (Fig. 6a-c), the arousal-relevant OA neurons sVUMlb5 and sVPMmd2 are almost sensory devoid (1 and 2 modalities, respectively) (Fig. 6d-f). The larval connectome, in its current state of tracing, did not show any connections from VNC sensory neurons (mechanosensory and nociceptive) to OA-neuron axons. We also identified pathways that involve ascending neurons projecting this sensory information to the brain; the OA-neurons were the fifth or sixth order neuron downstream of VNC sensory neurons, which we did not consider as significant input. Thus, OA neurons driving arousal in early life are relatively disconnected from the brain itself and from the external environment.

**Fig. 6.**
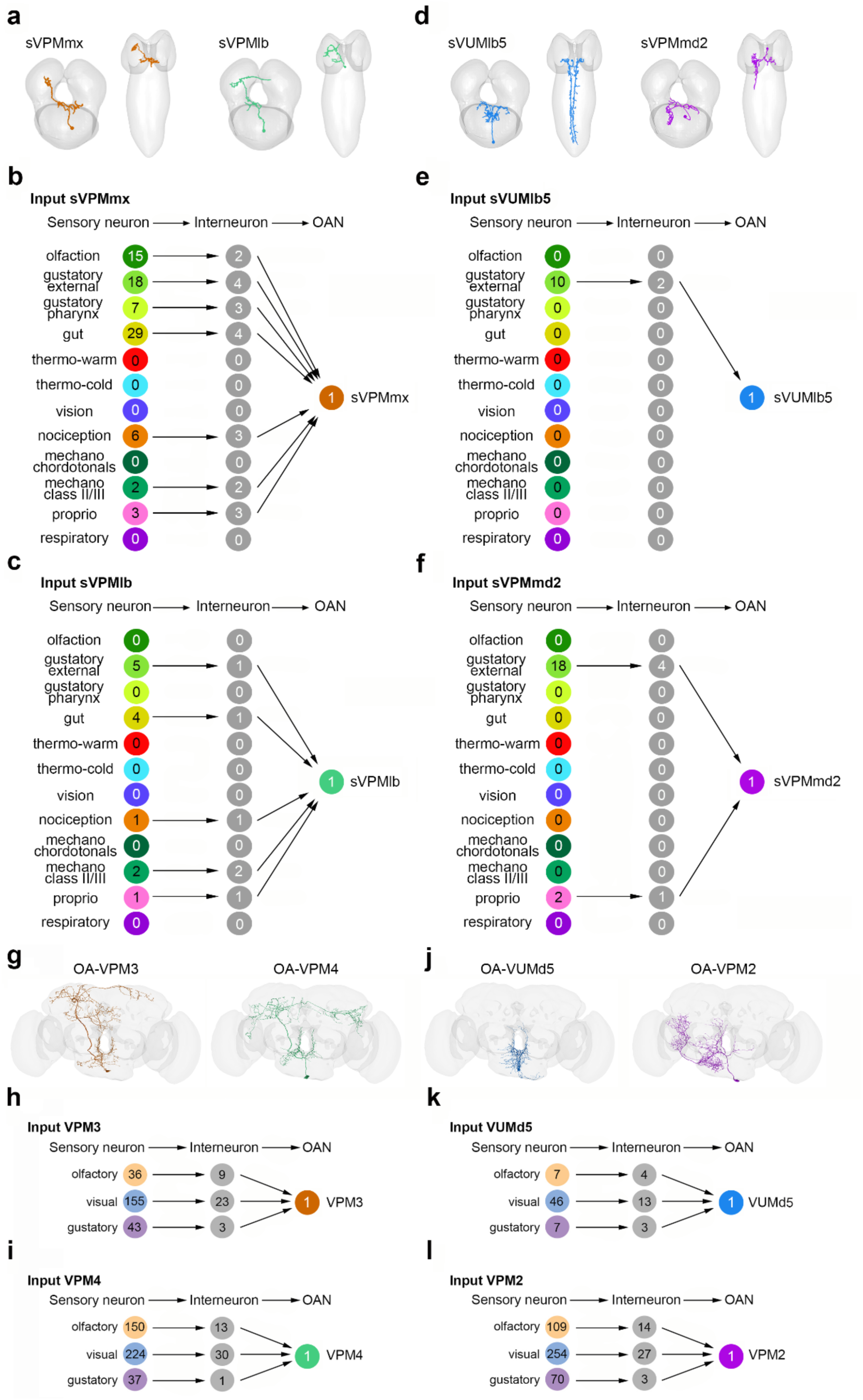
Connectome analysis reveals a developmental switch in sensory innervation of arousal-relevant OANs. **a,** Anterior and dorsal views of larval sVPMmx and sVPMlb, which do not promote arousal in larvae. **b,c,** Sensory modality inputs to sVPMmx and sVPMlb are extensive. **d,** Anterior and dorsal views of larval sVUMlb5 and sVPMmd2, which promote larval arousal. **e,f,** Sensory modality inputs to sVUMlb5 and sVPMmd2 are sparse. **g,** Anterior view of VPM3 and VPM4 neurons in the adult brain, which promote arousal at this life stage. **h,i,** Sensory input to VPM3 and VPM4 is extensive. **j,** Anterior view of VUMd5 and VPM2 neurons in the adult brain, which do not promote arousal at this stage. **k,l,** Sensory input to VUMd5 and VPM2.

We then performed a similar analysis of the adult OA SEZ neurons, in this case generating the specific number of sensory neurons (olfactory, visual, or gustatory) that are connected to a given OA cell. In contrast to larval stages, arousal-relevant OA neurons (OA-VPM3 and OA-VPM4) projected within the brain and received extensive sensory input, from 234 and 411 sensory neurons, respectively (Fig. 6g-i). OA-VUMd5, which was not sufficient to promote arousal in adulthood, had limited synaptic contacts in the brain and received far less sensory input, from only 60 cells. OA-VPM2 was also not sufficient to promote arousal in adulthood, but was unique in receiving little sensory input in the larval nervous system but dense sensory input in adulthood (from 433 neurons, Fig. 6j-l). Together, these findings suggest distinct circuit architectures promoting arousal across the lifespan, with larval arousal OA neurons insulated from direct sensory input and adult arousal OA neurons highly connected to sensory modalities.

## Discussion

Sleep is evident from the earliest developmental timepoints examined and continues until death. Yet, animals face dramatically different challenges and limitations across the lifespan that necessitate distinct sleep features. Primary drives in juvenile periods include growth and development, while repair and reproduction are prominent in maturity. Quantitative approaches support that sleep function transitions from neuronal reorganization to repair, and suggest such transitions are abrupt^66^, lending credence to the possibility that mechanistic switches could be required to generate functional change. In *Drosophila*, the neuromodulator octopamine plays a continuous role in promoting wakefulness from larval to adult periods^26,27^. Unexpectedly, we find that the specific OA neurons involved, their circuit architecture, and the nature of their sensory input are distinct in developmental compared to mature life stages. We propose that these differences reflect how sleep/wake circuits have evolved to meet the unique ecological demands facing an animal depending upon age.

Two classes of larval OA neurons (the paired sVPMmd2s and unpaired sVUMlb5) have been identified in this study as sufficient to suppress sleep and promote wakefulness. These neurons primarily project to the larval ventral nerve cord, an area most associated with processing central brain signals to coordinate locomotor output^67^. In contrast, the arousal-driving OA neurons of adulthood (OA-VPM3 and OA-VPM4) project extensively to brain regions involved in memory, sensory integration, and sleep itself. The distinct neurite morphologies potentially reflect distinct sleep functions. Larval OA arousal neurons are positioned to directly drive locomotor behavior, perhaps tightly coupling arousal to food-seeking exploration during this period of intensive growth. OA arousal-relevant neurons are swapped during metamorphosis, with the adult cells poised to influence higher cognitive functions like learning and memory. Connectomic analysis further refines this hypothesis by revealing larval sVPMmd2 and sVUMlb5 receive little direct sensory input, in contrast to adult OA-VPM3 and OA-VPM4. It is intriguing that gustatory input is one of the only represented sensory modalities among the larval OA arousal cells, consistent with the idea that a developing animal benefits from arousal systems that drive localizing nutrient sources. However, outside of feeding, other drives are subordinated to sleep, and thus arousal neurons are more disconnected from the surrounding sensory world to facilitate sleep unimpeded. In maturity, tight tuning to the external world is instead prioritized, reflected in dense sensory input to adult OA arousal neurons.

Might similar developmental switches in sleep/wake circuit logic occur in noradrenergic (NA) systems of the mammalian nervous system? NA neurons in the locus coeruleus (LC) of the mature mammalian brain promote arousal via widespread ascending projections^68–70^. LC neurons are activated by sensory stimuli^70,71^, consistent with a role for NA/OA in coupling arousal systems to sensory input in maturity across species. However, the LC also sends descending inputs to spinal cord, characterized in analgesia^72^, and innervates the parasympathetic nervous system to coordinate heart rate variations during sleep^73^. Findings in the fly should motivate exploration of such descending projections as potential mediators of NA-dependent arousal during gestation or in early postnatal periods.

Many behaviors in addition to sleep exhibit apparent continuity throughout development and into maturity, despite extensive nervous system structural remodeling during puberty in mammals and metamorphosis in insects. Feeding patterns change from developmental periods into adulthood but their behavioral expression is persistent and necessary for survival. Ingestive behaviors more broadly, however, present an interesting case study. In rats and other mammals, suckling and feeding are not developmentally continuous: suckling is not a necessary precursor to feeding, suckling is a transient state, and the two can exist in parallel^74^. By contrast, in *Drosophila*, crawling in limbless larva and walking in six-legged adults would seemingly share few commonalities aside from both being locomotor behaviors. Yet, the same pair of synaptically-connected interneurons generates the same locomotor behavior at both stages despite connectivity being lost and then re-established during metamorphosis^75^. There has been longstanding debate as to whether infant and mature sleep are truly continuous behaviors, or if juvenile sleep is an undifferentiated “pre-sleep” state that serves as precursor of adult forms (REM and NREM)^74,76^. Sleep is largely defined behaviorally in flies, and we observe a near total correspondence of early larval to adult sleep features^5,26,77^. An exception is the circadian control of sleep timing, evident in the adult but not in L2^26^. Recently, we pinpointed a cellular mechanism timed to the third instar larval stage (L3) in which a new synaptic connection forms between clock and arousal neurons, bringing sleep patterns under circadian control^78^. This mechanism supports a model in which foundational components of sleep are elaborated upon over developmental time, ultimately giving rise to mature forms; this elaboration model is consistent with temporally-sequenced emergence and coalescence of REM sleep features over the first two postnatal weeks in rats. Does developmental continuity of sleep necessitate shared underlying mechanisms? To the contrary, unique cellular and genetic mechanisms might govern sleep during specific epochs across the lifespan, and dedicated molecular cues likely guide the sleep maturation process itself^19^. Our findings support the continuous role of OA as a wake-promoting signal spanning larva to adulthood, but via dissociable OA neuronal subclasses with morphologies and sensory inputs distinctive to each life stage. We posit that sleep regulatory mechanisms change dynamically across developmental periods to support evolving sleep functions.

## Materials and methods

### Fly stocks

The following lines have been maintained as lab stocks: Iso31, UAS-TrpA1, and SS4268-Gal4. The GAL4 lines for the initial screen (R16B04, R57F09, R76G04, R76G06, R76G07, R76G08, R76G09, R76G10, R76G11, R76H01, R76H03, R76H04, R76H05, R76H07) the split-Gal4 lines R76H04-GAL4.DBD, Tdc2-p65.AD, MB113C-Gal4 and UAS-IVS-CsChrimson.mVenus were obtained from the Bloomington Drosophila Stock Center (BDSC). The trans-Tango MKII [MS1.1] line was obtained from Gilad Barnea [MK2.1] (BDSC #95317). The immortalization stock was a gift from Chris Doe: P[13XLexAop2-IVS-CsChrimson.mVenus]attP18; Actin5C-FRT-STOP-FRT-lexAop::65; pJFRC108-20XUAS-IVS-hPR::Flp-p10.

### Larval rearing and sleep assays

Adult flies were maintained on a standard molasses-based diet (8.0% molasses, 0.55% agar, 0.2% Tegosept, 0.5% propionic acid) at 25°C on a 12:12 light:dark (LD) cycle. To collect synchronized L2 larvae, adult flies were placed in an embryo collection cage (Genesee Scientific, catalog no. 59-100) and eggs were laid on a petri dish containing 3% agar, 2% sucrose, and 2.5% apple juice with yeast paste on top. Animals developed on this media for 2 days.

Sleep assays for L2 larvae were performed using the LarvaLodge multi-well device as described previously^26,79^. Briefly, molting L2 larvae were placed into individual wells of the LarvaLodge containing 90 μl of 3% agar and 2% sucrose media covered with a thin layer of yeast paste. The LarvaLodge was covered with a transparent acrylic sheet and placed into a DigiTherm (Tritech Research) incubator at 25°C for imaging. Experiments were performed in the dark. For thermogenetic experiments, animals were maintained at 22°C. Larvae were then placed into the LarvaLodge (as described above) which was moved into a DigiTherm (Tritech Research) incubator at 30°C for imaging.

### Immortalization and larval closed-loop optogenetic activation

Adult flies were allowed to lay eggs on apple juice plates in egg laying chambers. Freshly hatched first instar larvae were collected and moved onto a new plate and allowed to feed on 100 μl yeast paste containing 2 μl of 50 mM RU-486 (M8046, Sigma) for 24 hr. Larvae were allowed to grow into adults and the brains with the VNC attached were dissected out for standard staining and imaging procedure. For larval optogenetic activation of immortalized neurons, 1st instar larvae were collected and fed with 50 μl of 40 mM all trans-retinal (ATR, R2500, Sigma) and 2 μl of 50 mM RU-486 mixed with 100 μl yeast paste overnight. For imaging sessions the next day, molting L1 larvae were collected and placed into a LarvaLodge and the lodge placed into a fan-ventilated enclosure containing 830 nm IR LEDs providing dark field illumination (Fig. 2h). We acquired images every 6 seconds using a monochrome CMOS camera (Basler acA4024-29um) and monitored sleep as previously described^26,79^. A computer running custom Python software processed acquired images in real time and administered red photostimuli for 5 seconds to individual wells when a quiescent larva was detected (closed-loop) using a DLP-based projector (InFocus IN116xv).

### Adult thermogenetic and optogenetic activation and sleep assays

Animals were reared at 18°C to prevent activation of TrpA1 during development. Adult female flies were collected 2 to 3 days after eclosion and aged at 18°C on standard fly food on a 12-hour:12-hour LD cycle. Flies aged 5 to 7 days were anesthetized on CO2 pads (Genesee Scientific, catalog no. 59-114) and loaded into individual glass tubes (DAM tubes with 5% sucrose and 2% agar) for monitoring locomotor activity in MultiBeam Activity Monitors (Trikinetics, Waltham, MA). Data collection began at ZT0 at least 24 hours following CO2 anesthesia at 22°C on a 12-hour:12-hour LD schedule for 2 days to measure baseline sleep. Activity was measured in 1-min bins, and sleep was defined as 5 min of consolidated inactivity. TrpA1 activation was achieved by a temperature shift to 30°C during the 12-hour dark period. For optogenetic experiments, newly eclosed adult flies were entrained on a 12:12 light:dark schedule for 3 days, then placed in constant dark with 1 mM ATR mixed into food for another 3 days. Flies were loaded into DAM tubes supplemented with 1 mM ATR and placed into MultiBeam Activity Monitors under constant dark conditions. For optogenetic activation, ILT Incubator Light (Trikinetics) delivered a constant red illumination during the second night of the experiment for 4 hours. Sleep data was collected and plotted from the 4^th^ hour of activation at ZT21. The following day animals were dissected and stained for the Venus tag to visualize the labeled neurons in each brain.

### Immunohistochemistry and imaging

Larval, pupal and adult brains were dissected in phosphate-buffered saline (PBS) and fixed in 4% paraformaldehyde for 20 min at room temperature. Following 3×10 min washes in 0.5% PBS with Triton X-100 (PBST), brains were incubated with primary antibody at 4°C overnight. Following 3×10 min washes in PBST, brains were incubated with secondary antibody at 4°C overnight. Following 3×10 min washes in PBST, brains were washed in 50:50 PBST:Glycerol for 20 minutes and then mounted in VECTASHIELD (Vector Laboratories). Primary antibodies included Rabbit anti-GFP (1:200; A-11122, Thermo Fisher Scientific), Chicken anti-GFP (1:200; A-10262, Thermo Fisher Scientific), Rabbit anti-Tdc2 (1:500; ab128225, Abcam) and Rabbit anti-HA (1:500; #3724, cell Signaling Technology). Secondary antibodies included Alexa Fluor Goat anti-Rabbit 488 (1:250; A-32731, Thermo Fisher Scientific), Alexa Fluor Goat anti-Rabbit 594 (1:250; A-32740, Thermo Fisher Scientific) and Alexa Fluor Goat anti-Chicken 488 (1:250; A-32931, Thermo Fisher Scientific). Brains were imaged with a Leica SP8 confocal microscope.

### Transsynaptic circuit mapping

For labeling postsynaptic partners of OA-VPM3 (OASEZ-split) neurons in the adult brain we used trans-Tango MkII according to the authors description^48^.

### Larval connectomic analysis

We analyzed the published connectome of the larval brain^60^. First, we identified the eight larval OA-neurons by checking the annotations in CATMAID for this dataset. Only sVPMmx had been previously identified. We then manually searched for neurons consistent in morphology with our light level images and identified the remaining six larval neurons. The descending neurons were not fully reconstructed; we thus completed their reconstruction where necessary. We used the published^60^ annotations of sensory neurons and checked for direct connections to OA-neurons in CATMAIDs graph widget, which did not exist. We thus searched for indirect connections using the “grow” tool to identify interneurons downstream of the sensory neurons and upstream of OA-neurons. We then summed the connections across the two hemispheres and counted the interneurons participating in those connections. The “grow” tool was also used to search for longer paths (more than two hops) from sensory to OA-neurons.

### Adult connectomic analysis

Codex (FlyWire Brain Dataset FAFB v783) and FlyWire Neuroglancer were used to identify and visualize OA cell types within an adult female brain EM dataset^56,57,64,65^ based on the single-cell descriptions^44^.

FlyWire IDs: 720575940623503318 (OA-VPM2), 720575940644745120 (OA-VPM3), 720575940625264457 (OA-VPM4), 720575940630956345 (OA-VUMd3), and 720575940625489160 (OA-VUMd5).

Sensory inputs to each OAN were identified using Codex (v783, January 2025)^57,61–63,65^. All connections were thresholded at 5 synapses. The sensory neurons were clustered into three modalities:

(i) **olfactory** - antennal nerve cells (AN), antennal lobe projection neurons (ALPN) and antennal lobe output neurons (ALON); (ii) **visual** - visual_projection and OL_intrinsic; and (iii) **gustatory** (taxonomy^61^).

### Statistical Analysis

All statistical analysis was done in GraphPad (Prism). For comparisons between two conditions, two-tailed unpaired t-tests were used. For comparisons between multiple groups, ordinary one-way ANOVAs followed by Tukey’s multiple comparison tests were used.

## Acknowledgments

We thank members of the Kayser Lab for helpful discussions and input. We thank the Princeton FlyWire team and members of the Murthy and Seung labs, as well as members of the Allen Institute for Brain Science, for development and maintenance of FlyWire (supported by BRAIN Initiative grants MH117815 and NS126935 to Murthy and Seung). We also acknowledge members of the Princeton FlyWire team and the FlyWire consortium for neuron proofreading and annotation.

## Funding

NIH DP2NS111996 (MSK)

NIH R35NS137329 (MSK)

Autism Spectrum Program of Excellence (MSK; Research gift to the University of Pennsylvania; Daniel J. Rader, principal investigator).

Burroughs Wellcome Career Award for Medical Scientists (MSK)

Deutsche Forschungsgemeinschaft, 441181781 (AT)

Deutsche Forschungsgemeinschaft, 426722269 (AT)

Deutsche Forschungsgemeinschaft, 432195391 (AT)

ESF Plus, 100649752 (AT)

NIH R35GM119844 (KA)

## Author contributions

Conceptualization: MS, MS, AST, MSK

Investigation: All authors

Writing – Original Draft: MS, MSK

Writing – Review and Editing: All authors

ProjectSupervision and Funding: MSK

## Data and materials availability

All data needed to evaluate the conclusions in the paper are present in the paper and/or the Supplementary Materials.

## Figure and Figure Legends

**Extended Data Fig. 1.**
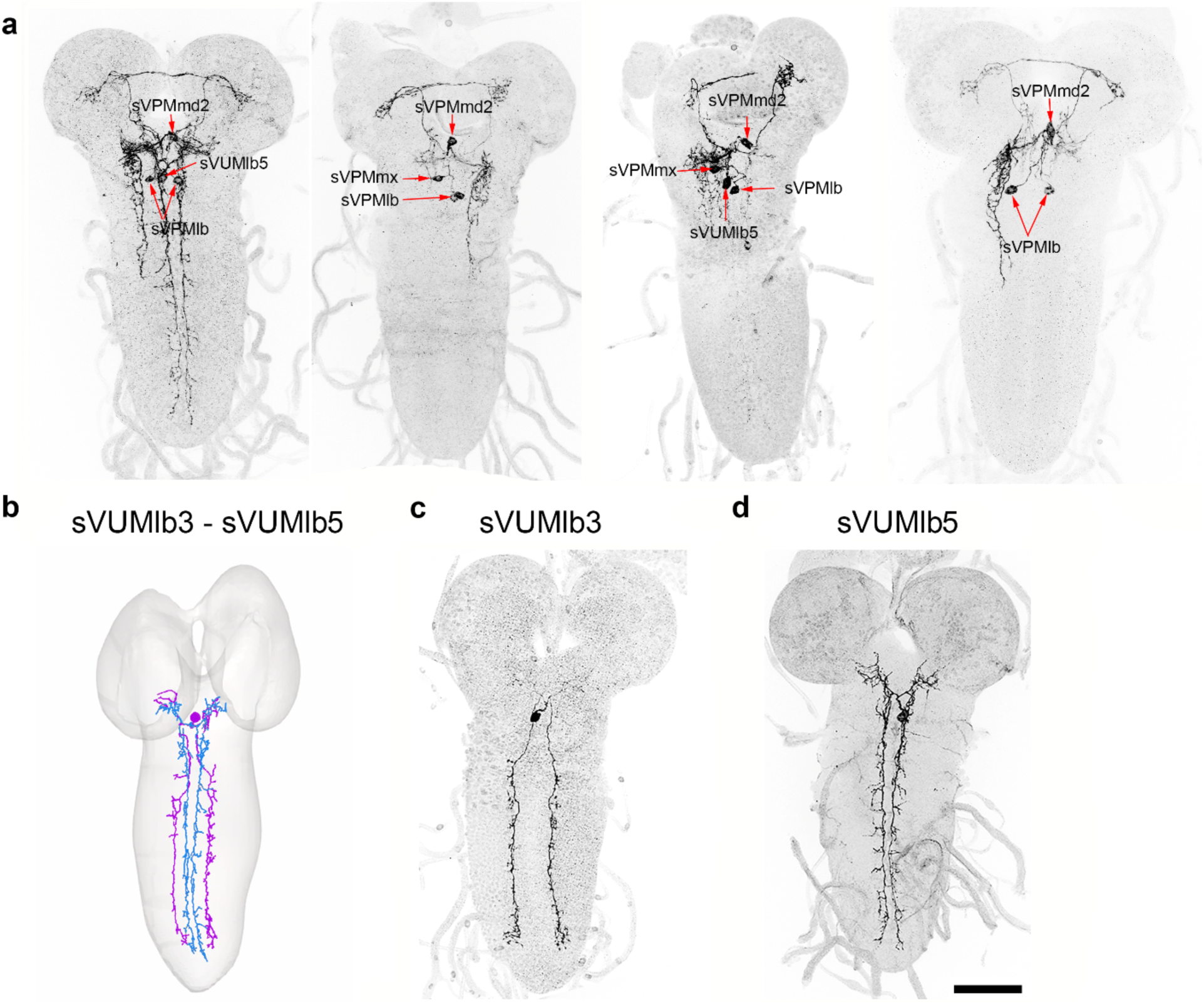
OA^SEZ^-split-GAL4 expression in larval nervous system is stochastic. **a,** 2nd instar larval brains with subsets of OANs labeled by OA^SEZ^-split-GAL4. Red arrows indicate cell bodies. The labeled neurons are sVPMlb, sVPMmx, sVPMmd2, sVUMlb3 and sVUMlb5. **b,** Reconstruction of the sVUMlb3 (purple) and sVUMlb5 (blue) neurons from the L1 larval connectome. **c,d,** Example images of sVUMlb3 and sVUMlb5 in L2 larval nervous systems. Scale bar: 50 µm

**Extended Data Fig. 2.**
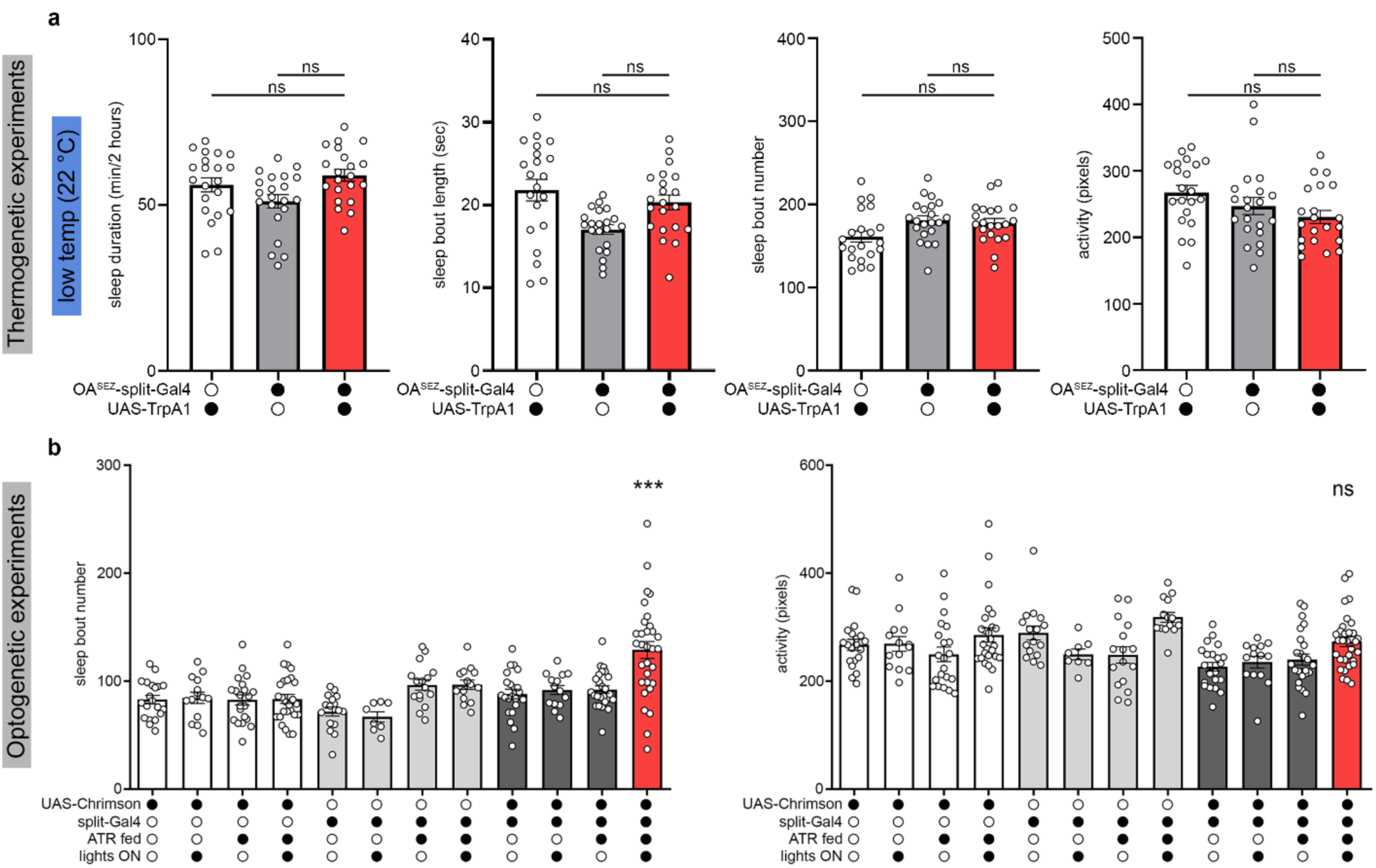
Additional controls and sleep metrics for larval OA^SEZ^-split-GAL4 thermo- and opto-genetic experiments. **a,** OA^SEZ^-split-GAL4 has no effect on sleep properties at restrictive temperatures (22 °C) compared to genetic controls (n=21). **b,** Optogenetic activation of the OA^SEZ^-split-GAL4 neurons in L2 increased sleep bout number without affecting activity during periods of wake.

**Extended Data Fig. 3.**
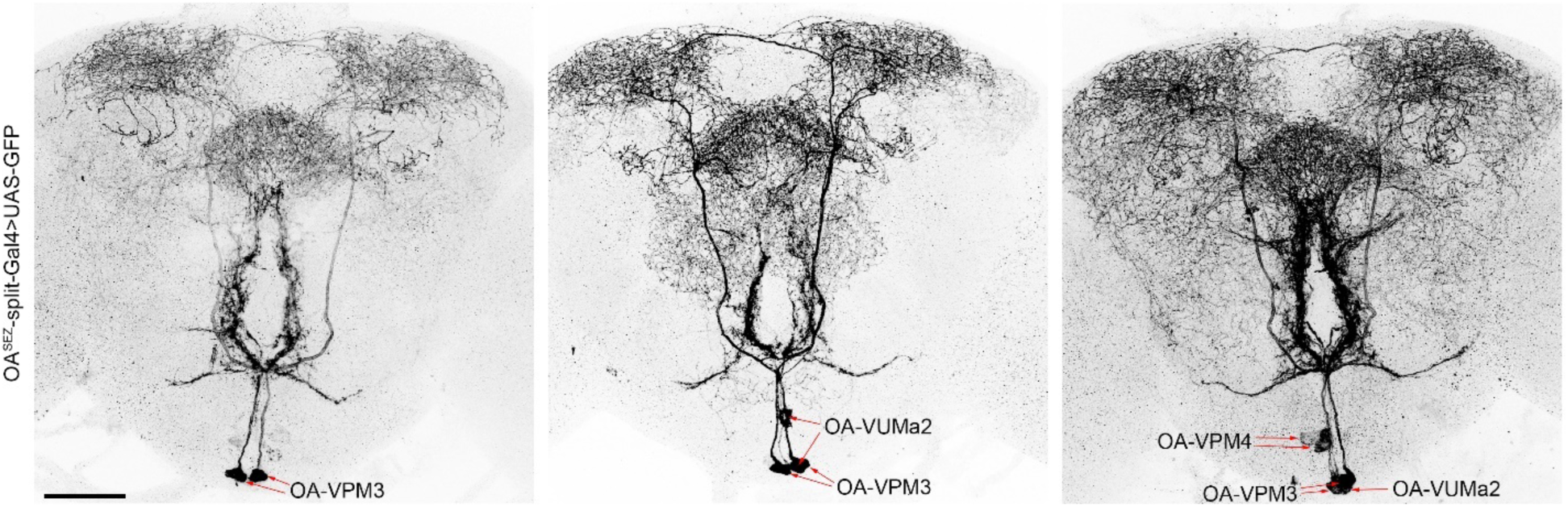
OA^SEZ^-split-GAL4 expression in adult brains is stochastic. Adult brains with subpopulations of OANs labeled by OA^SEZ^-split-GAL4. Red arrows indicate cell bodies. OA-VPM3 neurons are always present, and OA-VPM4s and OA-VUMa2s appear less commonly. Scale bar = 50 µm.

**Extended Data Fig. 4.**
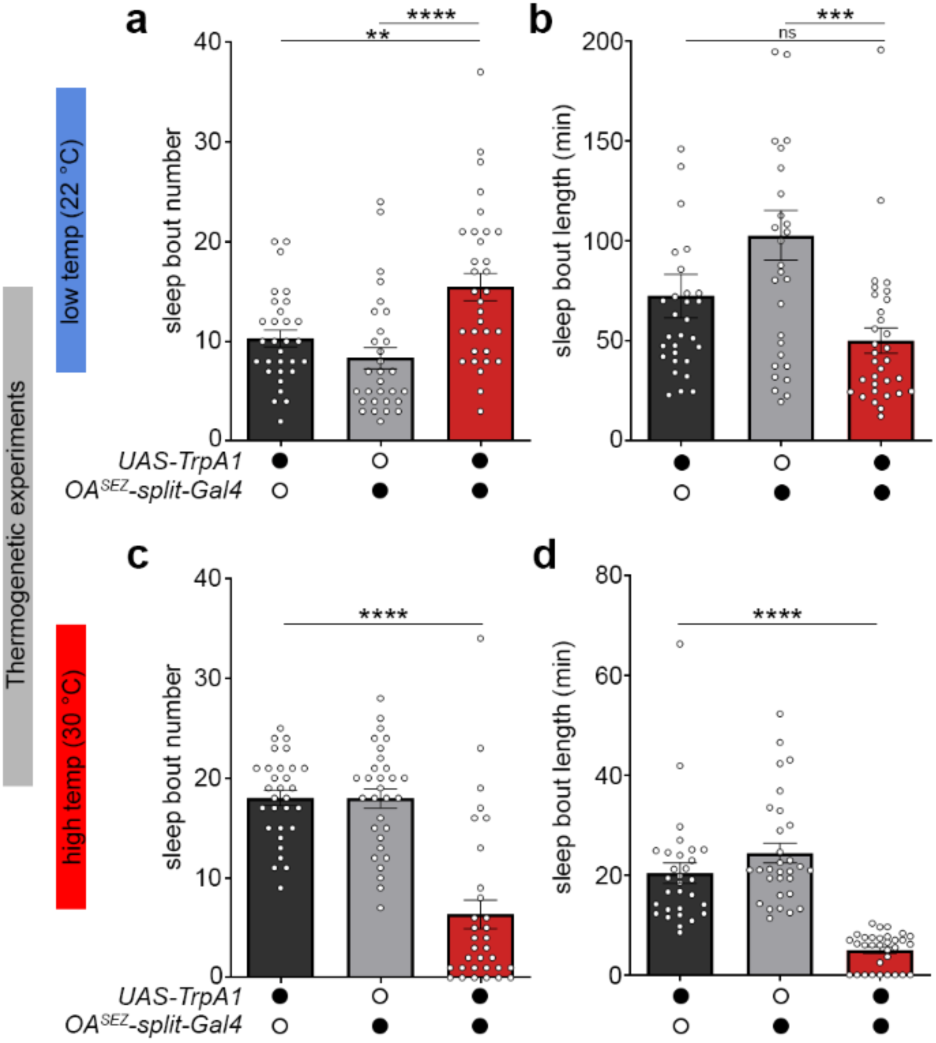
Additional sleep metrics for adult thermogenetic sleep experiment. **a,b,** Flies expressing UAS-TrpA1 under control of OA^SEZ^-split-GAL4 do not show reduced sleep bouts at restrictive temperatures (22 °C). **c,d,** At permissive temperatures (30 °C) both sleep bout number and duration are reduced (n=30-32).

**Extended Data Fig. 5.**
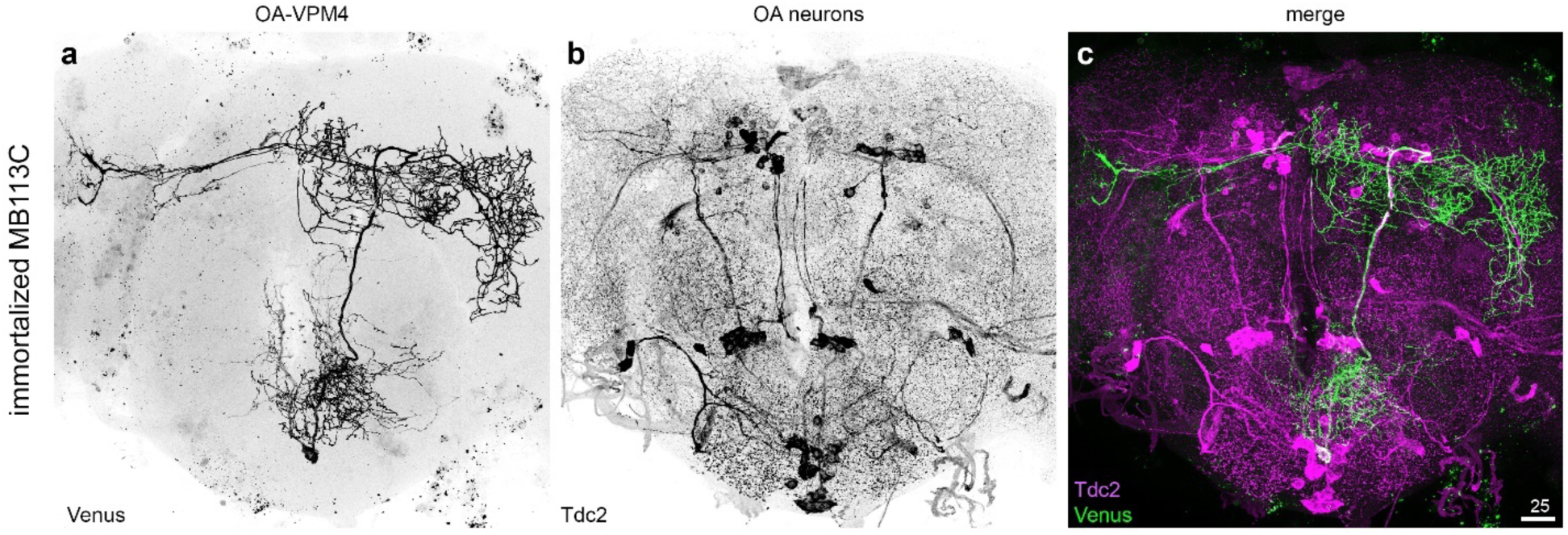
Larval OAN sVPMlb becomes OA-VPM4 in the adult brain. **a,** Immortalizing fluorescence in sVPMlb neurons during the 2nd instar stage using MB113C split-Gal4 and tracing into adulthood reveals larval sVPMlb becomes adult OA-VPM4. **b,** Visualization of adult OANs expressing Tdc2, with the merge image in **c**. Scale bar = 25 µm.

**Extended Data Fig. 6.**
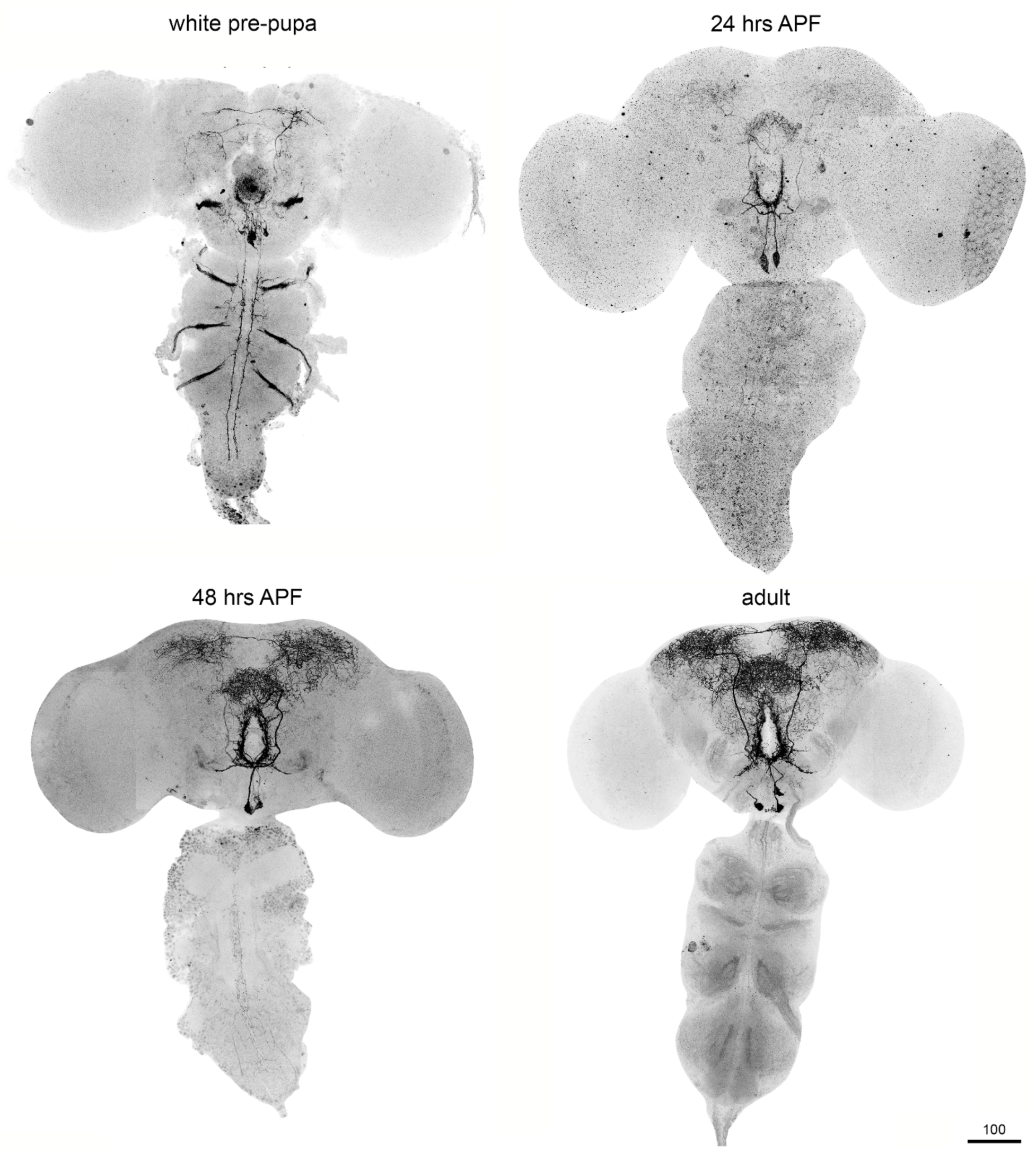
OA^SEZ^-split-GAL4 expression throughout metamorphosis. OA^SEZ^-split-GAL4 continues to label OANs through metamorphosis, though expression in descending neurons is lost after the white-prepupa stage. OA-VPM3 neurons are labeled most prominently during metamorphosis and in adulthood.

**Extended Data Fig. 7.**
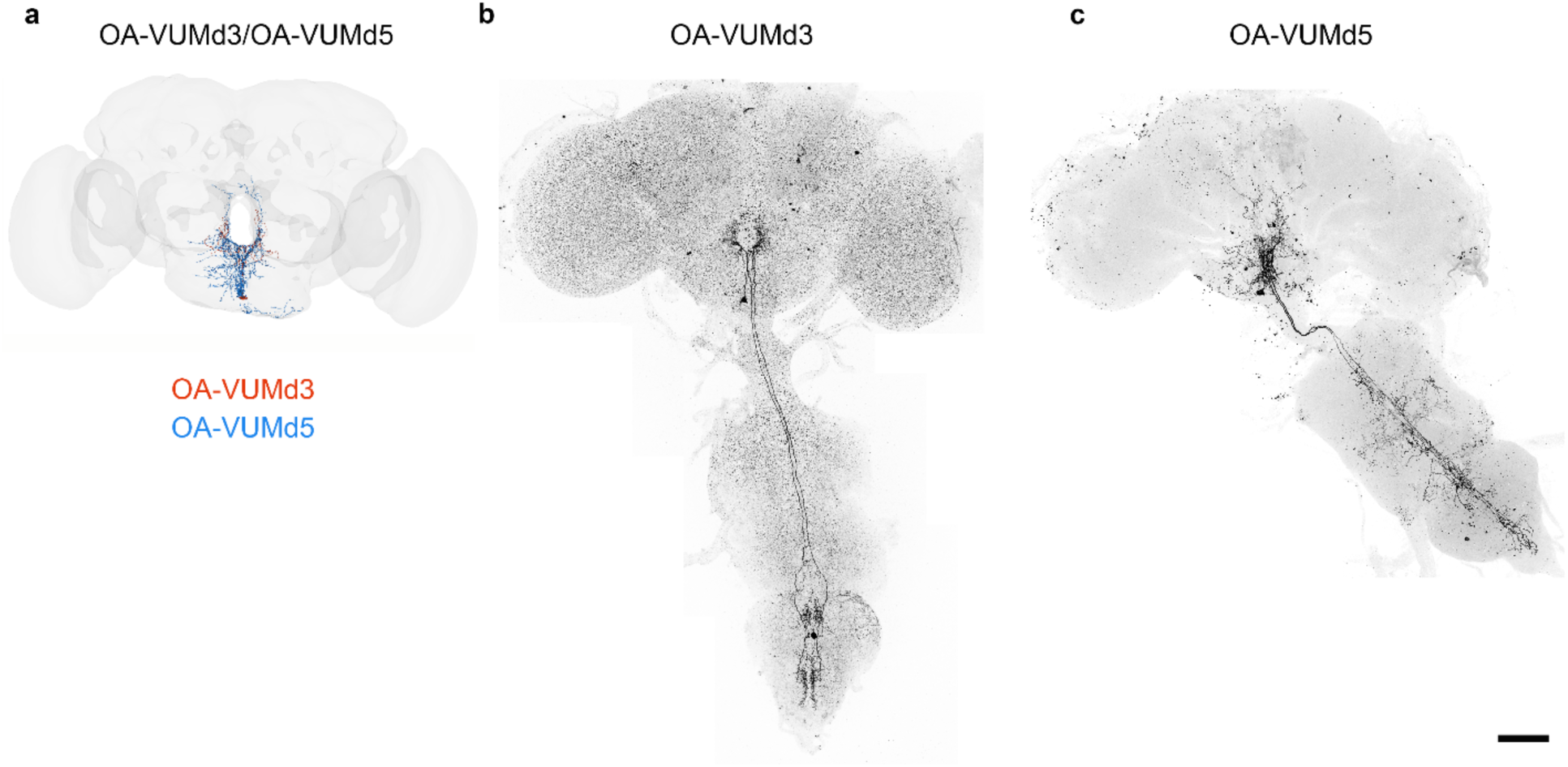
Distinct projection patterns of the unpaired OA neurons in the adult CNS. **a**, Reconstruction of the central brain projections of the OA-VUMd3 and OA-VUMd5 neurons from the adult brain connectome. **b**, A complete view of the OA-VUMd3 neuron including the VNC projections. **c**, A full reconstruction of the OA-VUMd5 neuron in the adult brain including the VNC. The images on **b** and **c** are combinations of multiple confocal stacks merged. Scale bar: **b**,**c** 100 µm.

**Extended Data Fig. 8.**
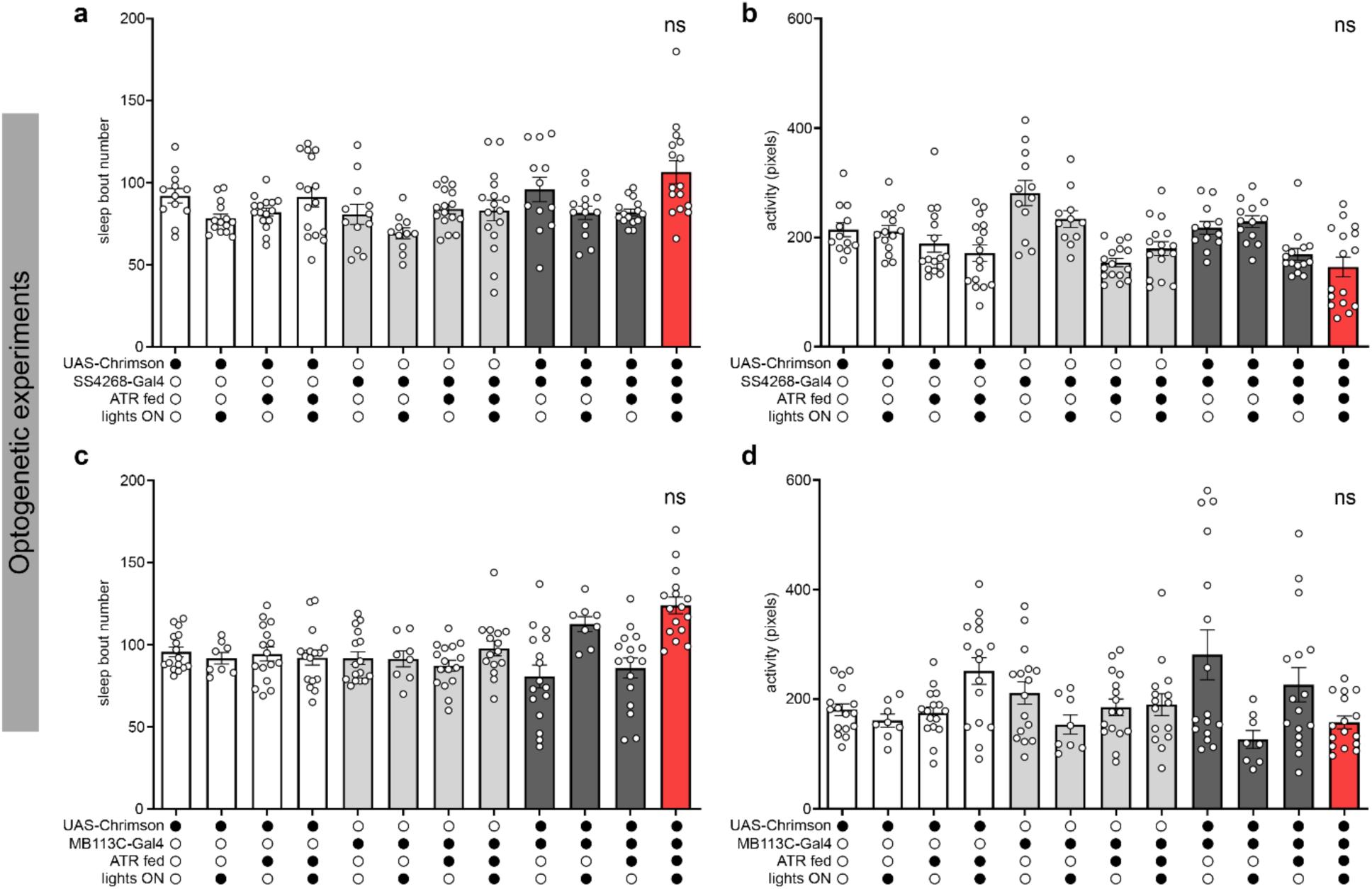
Additional sleep metrics related to activation of OA-sVPMmx and OA-sVPMlb. **a-d,** Optogenetic activation of the larval OA-sVPMmx or OA-sVPMlb did not affect sleep bout number or wake activity (n=8-16).

**Extended Data Fig. 9.**
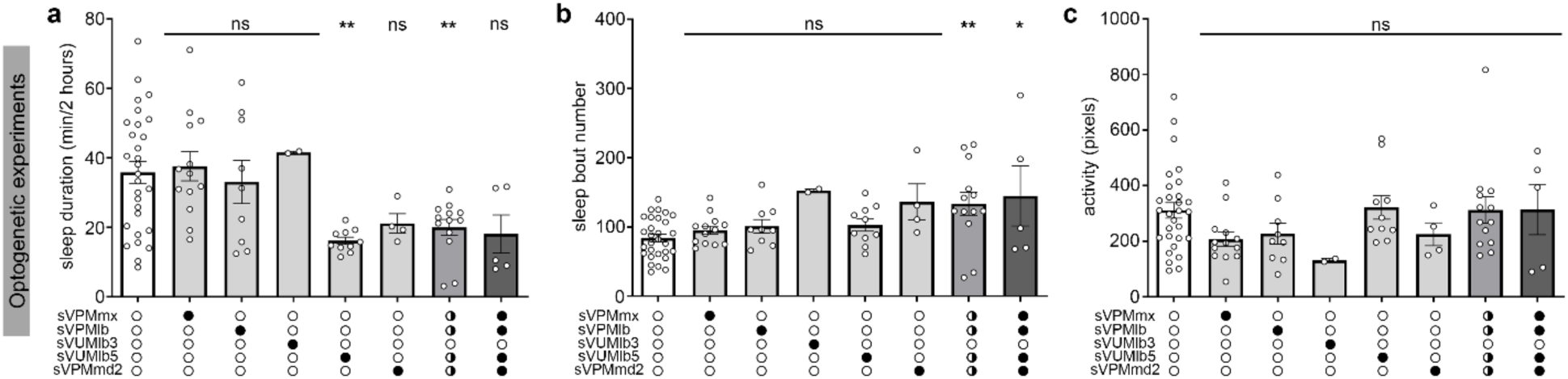
Additional sleep metrics related to stochastic labeling and activation of OA^SEZ^-split-GAL4 neurons. **a-c,** Sleep duration, sleep bout number, and activity data from stochastic labeling and optogenetic activation experiments using OA^SEZ^-split-GAL4 in 2nd instar larvae.

**Extended Data Fig. 10.**
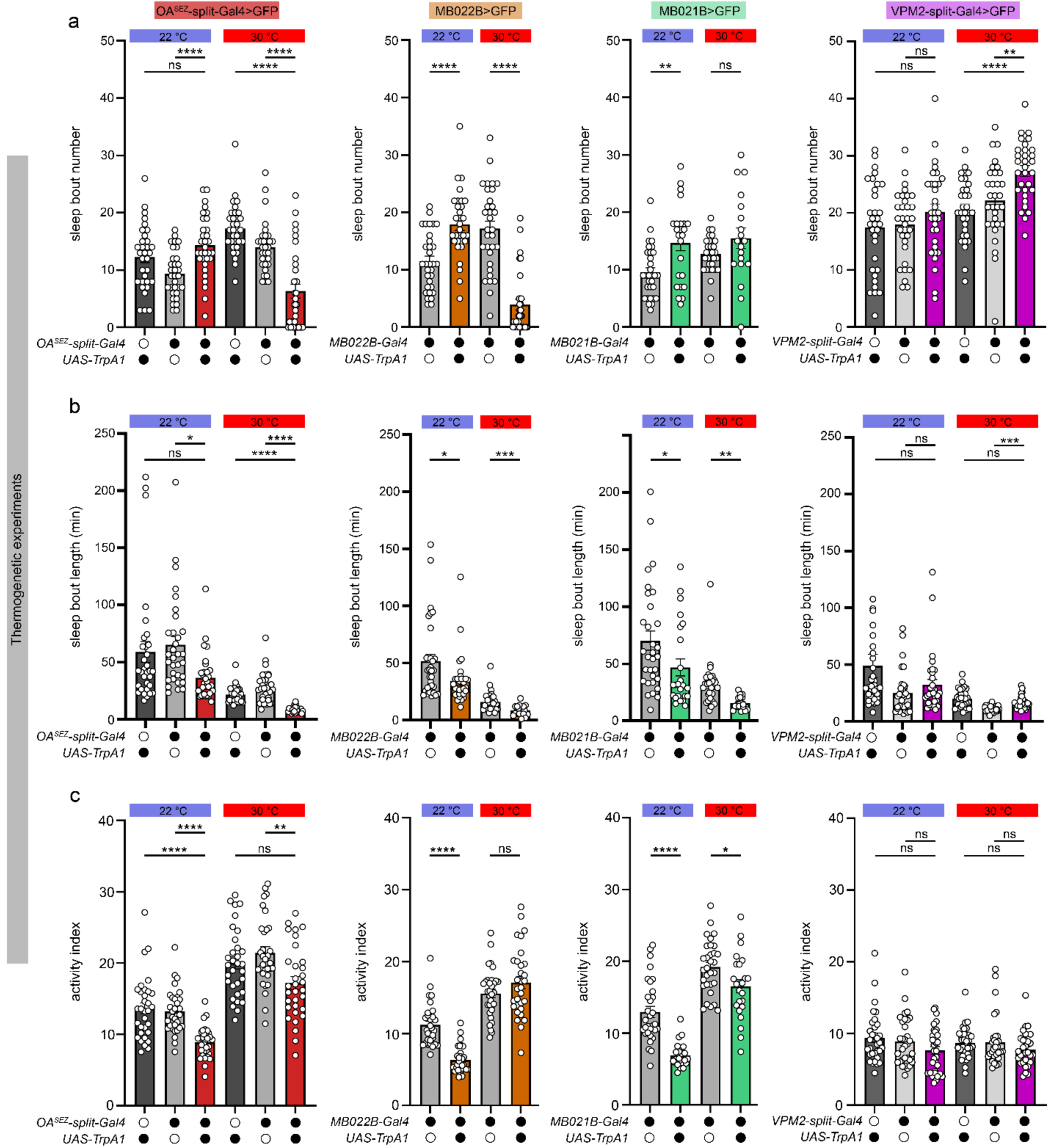
Additional sleep metrics related to adult OAN subpopulation activation in Figure 5. **a,** Sleep bout number, **b,** sleep bout length, and **c,** waking activity during thermogenetic activation of SEZ OAN subpopulations in adulthood. n=24-32.

**Extended Data Fig. 11.**
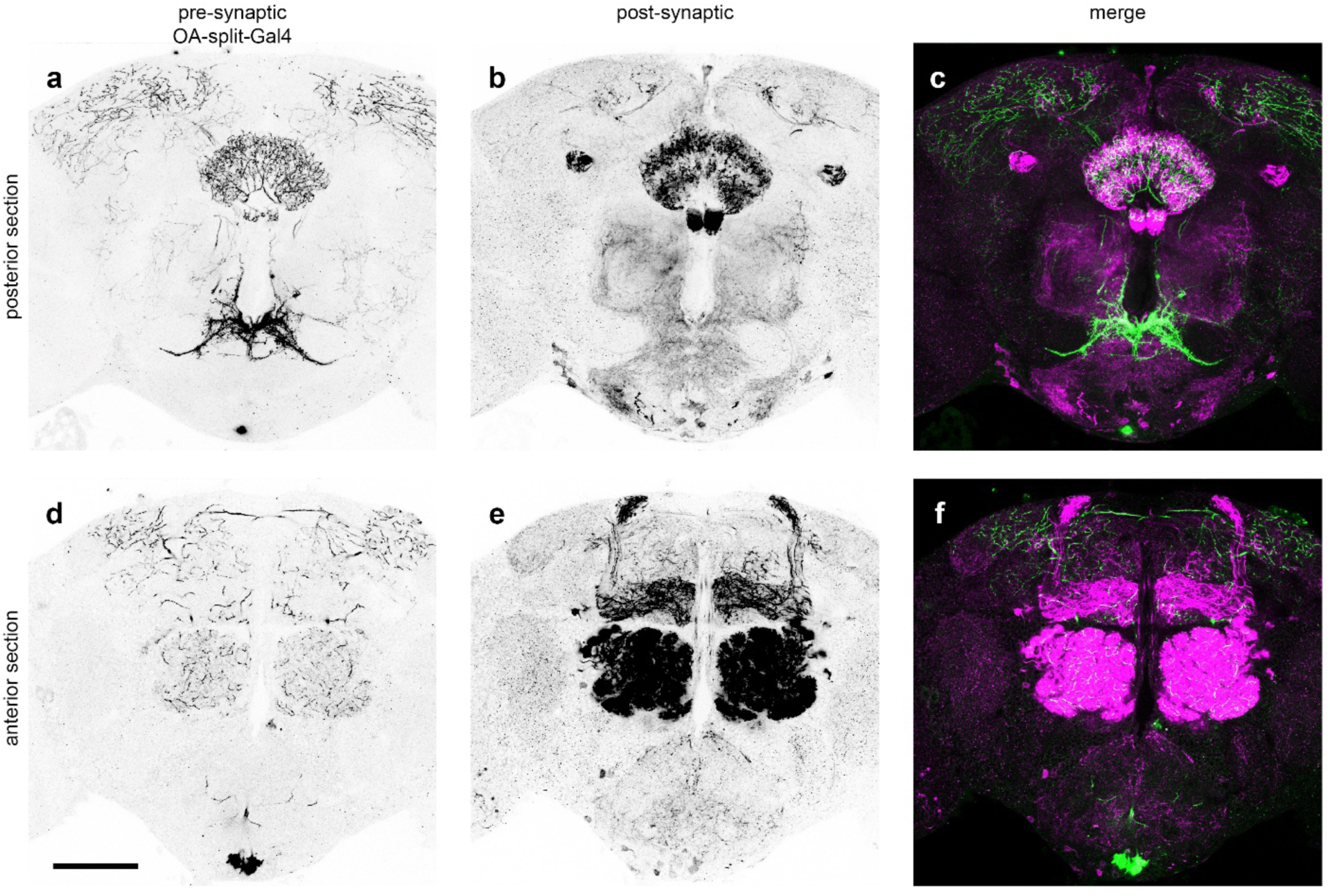
Trans-synaptic labeling of downstream partners of adult OA^SEZ^-split-GAL4 neurons. Posterior section of an adult brain labeling VPM3 projections (**a**) and their post-synaptic partners in the fan-shaped body (**b**). Merged image in **c**. Anterior section of the same brain show VPM3 projections (**d**) and downstream partners in the mushroom body and olfactory glomeruli (**e**). Merged image in **f**. Scale bar = 75 µm.

## References

1. Roffwarg, H. P., Muzio, J. N. & Dement, W. C. Ontogenetic development of the human sleep-dream cycle. Science 152, 604–19 (1966).

2. Kayser, M. S. & Biron, D. Sleep and Development in Genetically Tractable Model Organisms. Genetics 203, 21–33 (2016).

3. Blumberg, M. S., Dooley, J. C. & Tiriac, A. Sleep, plasticity, and sensory neurodevelopment. Neuron 110, 3230–3242 (2022).

4. Frank, M. G. Sleep and developmental plasticity not just for kids. Prog. Brain Res. 193, 221–232 (2011).

5. Shaw, P. J., Cirelli, C., Greenspan, R. J. & Tononi, G. Correlates of sleep and waking in Drosophila melanogaster. Science 287, 1834–7 (2000).

6. Kayser, M. S., Yue, Z. & Sehgal, A. A critical period of sleep for development of courtship circuitry and behavior in Drosophila. Science 344, 269–74 (2014).

7. Blumberg, M. S., Lesku, J. A., Libourel, P.-A., Schmidt, M. H. & Rattenborg, N. C. What Is REM Sleep? Curr. Biol. CB 30, R38–R49 (2020).

8. Gong, N. N. et al. Intrinsic maturation of sleep output neurons regulates sleep ontogeny in Drosophila. Curr. Biol. 32, 4025–4039.e3 (2022).

9. Dilley, L. C., Vigderman, A., Williams, C. E. & Kayser, M. S. Behavioral and genetic features of sleep ontogeny in Drosophila. Sleep 41, (2018).

10. Blumberg, M. S., Gall, A. J. & Todd, W. D. The development of sleep-wake rhythms and the search for elemental circuits in the infant brain. Behav Neurosci 128, 250–63 (2014).

11. Ednick, M. et al. A review of the effects of sleep during the first year of life on cognitive, psychomotor, and temperament development. Sleep 32, 1449–58 (2009).

12. Halbower, A. C. et al. Childhood obstructive sleep apnea associates with neuropsychological deficits and neuronal brain injury. PLoS Med. 3, e301 (2006).

13. Bian, W.-J., Brewer, C. L., Kauer, J. A. & de Lecea, L. Adolescent sleep shapes social novelty preference in mice. Nat. Neurosci. 10.1038/s41593-022-01076-8 (2022) doi:10.1038/s41593-022-01076-8.

14. Jones, C. E. et al. Early-life sleep disruption increases parvalbumin in primary somatosensory cortex and impairs social bonding in prairie voles. Sci. Adv. 5, eaav5188 (2019).

15. Mirmiran, M., van de Poll, N. E., Corner, M. A., van Oyen, H. G. & Bour, H. L. Suppression of active sleep by chronic treatment with chlorimipramine during early postnatal development: effects upon adult sleep and behavior in the rat. Brain Res. 204, 129–46 (1981).

16. Frank, M. G., Issa, N. P. & Stryker, M. P. Sleep enhances plasticity in the developing visual cortex. Neuron 30, 275–87 (2001).

17. Seugnet, L., Suzuki, Y., Donlea, J. M., Gottschalk, L. & Shaw, P. J. Sleep deprivation during early-adult development results in long-lasting learning deficits in adult Drosophila. Sleep 34, 137–46 (2011).

18. Dashti, H. S. et al. Genome-wide association study identifies genetic loci for self-reported habitual sleep duration supported by accelerometer-derived estimates. Nat. Commun. 10, 1100 (2019).

19. Chakravarti Dilley, L., et al. Identification of a Molecular Basis for the Juvenile Sleep State. eLife 9, e52676, (2020).

20. Koh, K. et al. Identification of SLEEPLESS, a sleep-promoting factor. Science 321, 372–376 (2008).

21. Cirelli, C. et al. Reduced sleep in Drosophila Shaker mutants. Nature 434, 1087–1092 (2005).

22. Kume, K., Kume, S., Park, S. K., Hirsh, J. & Jackson, F. R. Dopamine is a regulator of arousal in the fruit fly. J. Neurosci. Off. J. Soc. Neurosci. 25, 7377–7384 (2005).

23. Stavropoulos, N. & Young, M. W. insomniac and Cullin-3 Regulate Sleep and Wakefulness in Drosophila. Neuron 72, 964–976 (2011).

24. Weber, F. & Dan, Y. Circuit-based interrogation of sleep control. Nature 538, 51–59 (2016).

25. Shafer, O. T. & Keene, A. C. The Regulation of Drosophila Sleep. Curr. Biol. CB 31, R38– R49 (2021).

26. Szuperak, M. et al. A sleep state in Drosophila larvae required for neural stem cell proliferation. eLife 7, (2018).

27. Crocker, A., Shahidullah, M., Levitan, I. B. & Sehgal, A. Identification of a Neural Circuit that Underlies the Effects of Octopamine on Sleep:Wake Behavior. Neuron 65, 670–681 (2010).

28. Liu, Q., Liu, S., Kodama, L., Driscoll, M. R. & Wu, M. N. Two Dopaminergic Neurons Signal to the Dorsal Fan-Shaped Body to Promote Wakefulness in Drosophila. Curr. Biol. 22, 2114–2123 (2012).

29. Ueno, T. et al. Identification of a dopamine pathway that regulates sleep and arousal in Drosophila. Nat. Neurosci. 15, 1516–1523 (2012).

30. Selcho, M., Pauls, D., El Jundi, B., Stocker, R. F. & Thum, A. S. The role of octopamine and tyramine in Drosophila larval locomotion. J. Comp. Neurol. 520, 3764–3785 (2012).

31. Selcho, M., Pauls, D., Huser, A., Stocker, R. F. & Thum, A. S. Characterization of the octopaminergic and tyraminergic neurons in the central brain of Drosophila larvae. J. Comp. Neurol. 522, 3485–3500 (2014).

32. Jenett, A. et al. A GAL4-Driver Line Resource for Drosophila Neurobiology. Cell Rep. 2, 991–1001 (2012).

33. Hamada, F. N. et al. An internal thermal sensor controlling temperature preference in Drosophila. Nature 454, 217–220 (2008).

34. Dionne, H., Hibbard, K. L., Cavallaro, A., Kao, J. C. & Rubin, G. M. Genetic Reagents for Making Split-GAL4 Lines in Drosophila. Genetics 209, 31–35 (2018).

35. Saumweber, T. et al. Functional architecture of reward learning in mushroom body extrinsic neurons of larval Drosophila. Nat. Commun. 9, 1104 (2018).

36. Werkhoven, Z. et al. The structure of behavioral variation within a genotype. eLife 10, e64988 (2021).

37. Crocker, A. & Sehgal, A. Octopamine Regulates Sleep in Drosophila through Protein Kinase A-Dependent Mechanisms. J. Neurosci. 28, 9377–9385 (2008).

38. Machado, D. R. et al. Identification of octopaminergic neurons that modulate sleep suppression by male sex drive. eLife 6, e23130 (2017).

39. Truman, J. W., Price, J., Miyares, R. L. & Lee, T. Metamorphosis of memory circuits in Drosophila reveals a strategy for evolving a larval brain. eLife 12, e80594 (2023).

40. Pfeiffenberger, C., Lear, B. C., Keegan, K. P. & Allada, R. Locomotor Activity Level Monitoring Using the Drosophila Activity Monitoring (DAM) System. Cold Spring Harb. Protoc. 2010, pdb.prot5518 (2010).

41. Carreira-Rosario, A. et al. MDN brain descending neurons coordinately activate backward and inhibit forward locomotion. eLife 7, e38554 (2018).

42. Li, H. H. et al. A GAL4 driver resource for developmental and behavioral studies on the larval CNS of Drosophila. Cell Rep 8, 897–908 (2014).

43. Babski, H., Codianni, M. & Bhandawat, V. Octopaminergic descending neurons in Drosophila: Connectivity, tonic activity and relation to locomotion. Heliyon 10, (2024).

44. Busch, S., Selcho, M., Ito, K. & Tanimoto, H. A map of octopaminergic neurons in the Drosophila brain. J. Comp. Neurol. 513, 643–667 (2009).

45. Eschbach, C. et al. Recurrent architecture for adaptive regulation of learning in the insect brain. Nat Neurosci 23, 544–555 (2020).

46. Reyes, M. et al. Octopamine regulates neural circuits in the mushroom body and central complex, influencing sleep and context-dependent arousal. 2025.03.06.641836 Preprint at 10.1101/2025.03.06.641836 (2025).

47. Artiushin, G. & Sehgal, A. The Drosophila Circuitry of Sleep–Wake Regulation. Curr. Opin. Neurobiol. 44, 243–250 (2017).

48. Sorkaç, A., Savva, Y. A., Savaş, D., Talay, M. & Barnea, G. Circuit analysis reveals a neural pathway for light avoidance in Drosophila larvae. Nat. Commun. 13, 5274 (2022).

49. Pitman, J. L., McGill, J. J., Keegan, K. P. & Allada, R. A dynamic role for the mushroom bodies in promoting sleep in Drosophila. Nature 441, 753–756 (2006).

50. Joiner, W. J., Crocker, A., White, B. H. & Sehgal, A. Sleep in Drosophila is regulated by adult mushroom bodies. Nature 441, 757–60 (2006).

51. Sitaraman, D. et al. Propagation of Homeostatic Sleep Signals by Segregated Synaptic Microcircuits of the Drosophila Mushroom Body. Curr. Biol. 25, 2915–2927 (2015).

52. Donlea, J. M., Thimgan, M. S., Suzuki, Y., Gottschalk, L. & Shaw, P. J. Inducing sleep by remote control facilitates memory consolidation in Drosophila. Science 332, 1571–1576 (2011).

53. Donlea, J. M. et al. Recurrent Circuitry for Balancing Sleep Need and Sleep. Neuron 97, 378–389 4 (2018).

54. Liu, S., Liu, Q., Tabuchi, M. & Wu, M. N. Sleep Drive Is Encoded by Neural Plastic Changes in a Dedicated Circuit. Cell 165, 1347–1360 (2016).

55. Aso, Y. et al. Mushroom body output neurons encode valence and guide memory-based action selection in Drosophila. eLife 3, e04580 (2014).

56. Dorkenwald, S. et al. FlyWire: online community for whole-brain connectomics. Nat. Methods 19, 119–128 (2022).

57. Dorkenwald, S. et al. Neuronal wiring diagram of an adult brain. Nature 634, 124–138 (2024).

58. Donlea, J. M., Pimentel, D. & Miesenböck, G. Neuronal machinery of sleep homeostasis in Drosophila. Neuron 81, 860–872 (2014).

59. Eichler, K. et al. The complete connectome of a learning and memory centre in an insect brain. Nature 548, 175–182 (2017).

60. Winding, M. et al. The connectome of an insect brain. Science 379, eadd9330 (2023).

61. Schlegel, P. et al. Whole-brain annotation and multi-connectome cell typing of Drosophila. Nature 634, 139–152 (2024).

62. Buhmann, J. et al. Automatic detection of synaptic partners in a whole-brain Drosophila electron microscopy data set. Nat. Methods 18, 771–774 (2021).

63. Heinrich, L., Funke, J., Pape, C., Nunez-Iglesias, J. & Saalfeld, S. Synaptic Cleft Segmentation in Non-isotropic Volume Electron Microscopy of the Complete Drosophila Brain. in Medical Image Computing and Computer Assisted Intervention – MICCAI 2018 (eds Frangi, A. F., Schnabel, J. A., Davatzikos, C., Alberola-López, C. & Fichtinger, G.) 317– 325 (Springer International Publishing, Cham, 2018). doi:10.1007/978-3-030-00934-2_36.

64. Matsliah, A., et al. Codex: Connectome Data Explorer. (2023). doi:10.13140/RG.2.2.35928.67844.

65. Zheng, Z. et al. A Complete Electron Microscopy Volume of the Brain of Adult Drosophila melanogaster. Cell 174, 730–743.e22 (2018).

66. Cao, J., Herman, A. B., West, G. B., Poe, G. & Savage, V. M. Unraveling why we sleep: Quantitative analysis reveals abrupt transition from neural reorganization to repair in early development. Sci. Adv. 6, eaba0398 (2020).

67. Hunter, I., Coulson, B., Zarin, A. A. & Baines, R. A. The Drosophila Larval Locomotor Circuit Provides a Model to Understand Neural Circuit Development and Function. Front. Neural Circuits 15, (2021).

68. Saper, C. B., Fuller, P. M., Pedersen, N. P., Lu, J. & Scammell, T. E. Sleep state switching. Neuron 68, 1023–1042 (2010).

69. Carter, M. E. et al. Tuning arousal with optogenetic modulation of locus coeruleus neurons. Nat. Neurosci. 13, 1526–1533 (2010).

70. Osorio-Forero, A., Cherrad, N., Banterle, L., Fernandez, L. M. J. & Lüthi, A. When the Locus Coeruleus Speaks Up in Sleep: Recent Insights, Emerging Perspectives. Int. J. Mol. Sci. 23, 5028 (2022).

71. Hayat, H. et al. Locus coeruleus norepinephrine activity mediates sensory-evoked awakenings from sleep. Sci. Adv. 6, eaaz4232 (2020).

72. Hirschberg, S., Li, Y., Randall, A., Kremer, E. J. & Pickering, A. E. Functional dichotomy in spinal-vs prefrontal-projecting locus coeruleus modules splits descending noradrenergic analgesia from ascending aversion and anxiety in rats. eLife 6, e29808 (2017).

73. Osorio-Forero, A. et al. Noradrenergic circuit control of non-REM sleep substates. Curr. Biol. 31, 5009–5023.e7 (2021).

74. Blumberg, M. S. HOMOLOGY, CORRESPONDENCE, AND CONTINUITY ACROSS DEVELOPMENT: THE CASE OF SLEEP. Dev. Psychobiol. 55, 92–100 (2013).

75. Lee, K. & Doe, C. Q. A locomotor neural circuit persists and functions similarly in larvae and adult Drosophila. eLife 10, e69767 (2021).

76. Frank, M. G. & Heller, H. C. The ontogeny of mammalian sleep: a reappraisal of alternative hypotheses. J. Sleep Res. 12, 25–34 (2003).

77. Hendricks, J. C. et al. Rest in Drosophila Is a Sleep-like State. Neuron 25, 129–138 (2000).

78. Poe, A. R. et al. Developmental emergence of sleep rhythms enables long-term memory capabilities in Drosophila. 2022.02.03.479025 Preprint at 10.1101/2022.02.03.479025 (2022).

79. Churgin, M. A. et al. Quantitative Imaging of Sleep Behavior in Caenorhabditis elegans and larval Drosophila melanogaster. Nat. Protoc. 14, 1455–1488 (2019).

